# Structural and functional characterization of polyclonal antibody responses reveals shared and distinct antigenic features of RSV and hMPV fusion proteins

**DOI:** 10.64898/2026.11.10.765330

**Authors:** Erick Bermúdez-Méndez, Sebastian Ols, Leonardo V. Cufaro, Shuhao Xiao, Zhitong Peng, Florian Gegenfurtner, Elise R. Eray, Marcos C. Miranda, Rodrigo Arcoverde Cerveira, Mikel Segura García, Miguel A. Véliz Pernilla, Leo Hanke, Andrew J. Borst, Neil P. King, Aleksandar Antanasijevic, Karin Loré

## Abstract

Respiratory syncytial virus (RSV) and human metapneumovirus (hMPV), members of the *Pneumoviridae* family, circulate widely and are leading causes of hospitalization for acute respiratory infections in infants and older adults. The viral fusion (F) protein in its prefusion (preF) conformation is the primary target of neutralizing antibodies, yet how naturally elicited circulating polyclonal antibodies are distributed across its antigenic landscape remains incompletely understood. Here, we combined serological profiling, site-specific antibody competition assays and electron microscopy-based polyclonal epitope mapping (EMPEM) to characterize preF- directed immunity to RSV and hMPV in healthy adults. All donors exhibited detectable preF- binding antibodies that positively correlated with virus neutralization and broadly recognized multiple antigenic sites, consistent with immunodominance patterns predicted by a surface-centric computational model. Despite this overall similarity, the distribution of responses across individual antigenic sites differed between RSV and hMPV. EMPEM analyses revealed consistent targeting of the highly neutralization-sensitive preF apex (site Ø) in both viruses, alongside less abundant responses against equatorial and upper-middle epitopes (sites II, III, IV and V). We additionally observed antibody-bound destabilized preF, supporting a model in which antibody binding promotes transient opening of the preF trimer and may interfere with the conformational rearrangements required for membrane fusion. Together, these findings highlight apex-directed antibodies as a prominent feature of naturally acquired pneumovirus immunity in adults and reveal both shared and distinct patterns of antigenic recognition that may inform next-generation vaccine design and antibody-based prophylactics.

## INTRODUCTION

Respiratory syncytial virus (RSV) and human metapneumovirus (hMPV), members of the *Pneumoviridae* family, are major causes of acute lower respiratory tract infections and represent a significant burden for public health^1,2^. Infants, older adults and immunocompromised individuals are particularly vulnerable, as infections often progress to severe disease requiring hospitalization and significantly increasing the risk of death^3–6^. These viruses circulate widely, with primary infections typically occurring in early childhood and subsequent exposures recurring throughout life^7,8^. In the case of RSV, vaccines have recently been licensed for older adults, adults at increased risk of severe disease and pregnant women, while prophylactic monoclonal antibodies (mAbs) can be administered to infants^9,10^. In contrast, no prophylactic or therapeutic measures specific to hMPV are currently approved.

Although effective protection against pneumoviruses likely involves both cellular and humoral immunity, the demonstrated efficacy of prophylactic RSV mAbs in preventing infection and RSV-associated hospitalization^11^ emphasizes the critical role of humoral responses. The viral fusion (F) protein is anchored on the virus surface and its role is to mediate entry by driving fusion of the viral and host cell membranes through a conformational transition from a prefusion (preF) to a postfusion (postF) state^12^. In its preF conformation, the F protein is the main target of neutralizing antibodies, from which seven distinct antigenic sites (Ø, I, II, III, IV, V and VI) have previously been described^13,14^.

Traditionally, pneumovirus antibody binding and neutralization have been measured using enzyme-linked immunosorbent assays (ELISAs) and virus neutralization assays, including in clinical settings^15–17^. While valuable and informative to confirm antigen-specific reactivity and neutralizing capacity, such methods do not directly provide information on the targeted epitopes. Conversely, traditional structural analyses have thus far only been performed using isolated mAbs, resulting in more than 50 pneumovirus F-mAb complex structures to date. However, these approaches do not address the complexity of the polyclonal antibody response, creating a disconnect between serology and structural analyses. Furthermore, isolated mAbs from memory B cell clones do not necessarily reflect the abundance of antibody specificities in serum and their contribution to humoral immunity^18,19^.

In recent years, electron microscopy-based polyclonal epitope mapping (EMPEM) has enabled the structural characterization of antibody specificities in polyclonal sera or plasma, providing new insights into humoral responses following infection or vaccination. To date, most EMPEM studies have focused on HIV Env^20–22^, influenza hemagglutinin^23,24^ and coronavirus spikes^25,26^, but no study has yet focused on pneumoviruses. Here, we characterized baseline humoral immunity to pneumoviruses in healthy adults using serological and EMPEM analyses of polyclonal responses to RSV and hMPV preF. This combined approach revealed detectable antibody reactivity across all main preF antigenic sites and consistent engagement of antigenic site Ø, a known epitope of neutralizing antibodies. Our findings provide the first comparative structural characterization of naturally acquired circulating antibody responses to RSV and hMPV, defining the baseline landscape of preF antigen recognition and revealing both common and virus-specific features of antibody targeting across pneumoviruses.

## RESULTS

### Circulating antibody responses to RSV and hMPV in healthy adults

We aimed to characterize the polyclonal antibody response to RSV and hMPV in healthy adults by combining conventional serological assays with EM-based techniques (**Fig. 1A**). Blood samples from 20 anonymous donors (D001-D020) were collected, and plasma was analyzed for preF- and postF-binding antibody levels, as well as virus-specific neutralizing titers. The pneumovirus F protein in its preF conformation is metastable and can spontaneously refold or respond to certain triggers, resulting in an irreversible transition from the preF to the postF form^13^. Because the integrity and conformation of the F proteins are critical for reliable interpretation of results, we performed both functional and structural validation of the RSV and hMPV preF proteins we utilized (**Fig. S1A-H**). PreF proteins exhibited the expected antigenicity profile, confirmed by binding of a virus-specific reference panel of mAbs known to target distinct antigenic sites (**Fig. S1C, G**). Negative-stain EM analysis further confirmed the proteins were homogeneous in their preF conformation and that previously reported atomic structures fitted well into the 3D maps (**Fig. S1D, H**).

**Figure 1.**
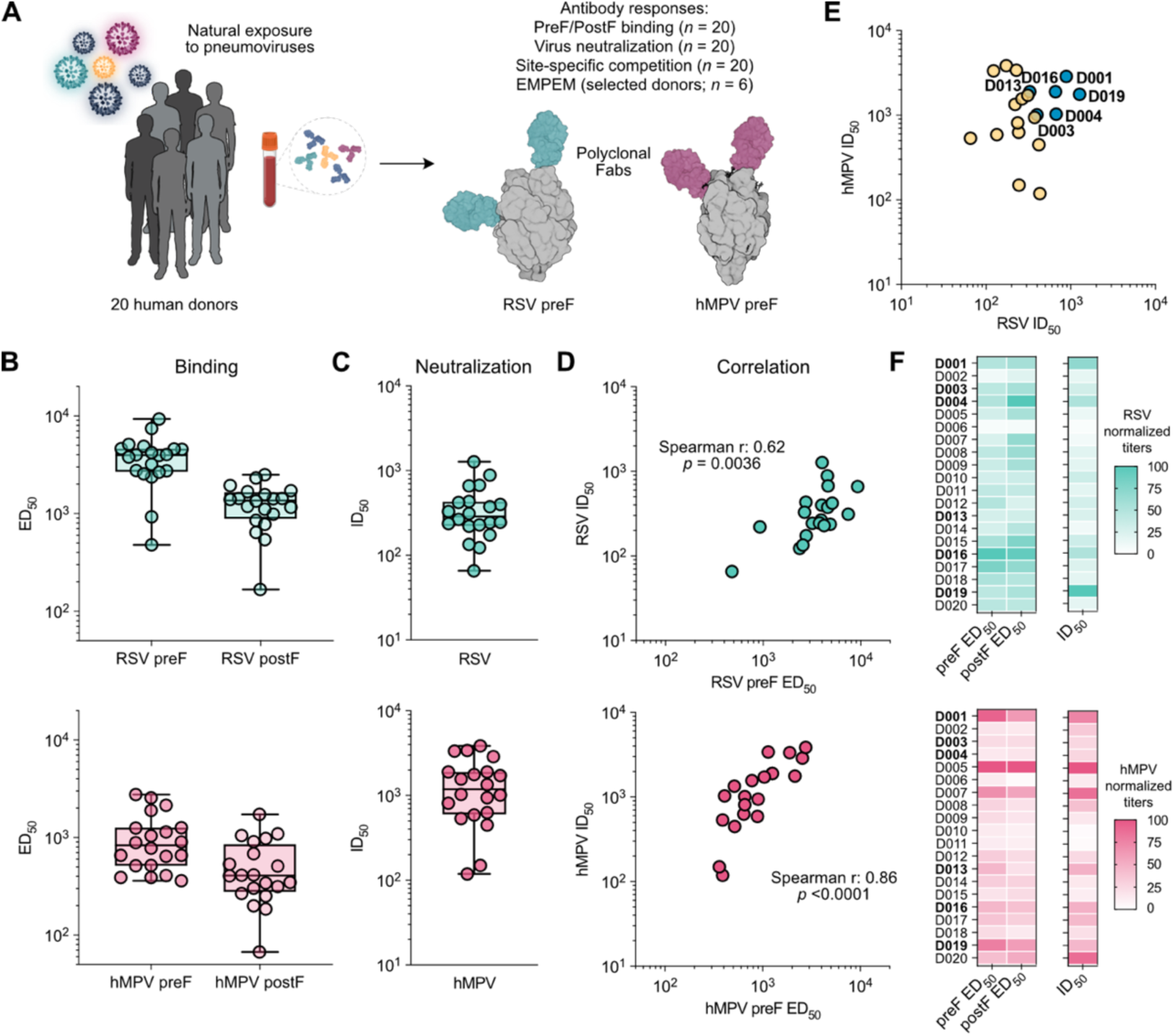
Circulating antibody responses to RSV and hMPV in healthy adults. A. Schematic illustration of the study design. Blood samples from 20 healthy human donors were collected and screened for their circulating antibodies against RSV and hMPV in a range of serological assays. B. RSV and hMPV prefusion (preF)- and postfusion (postF)-specific plasma IgG binding titers determined by ELISA. Graphs show half-maximal effective dilutions (ED50). Horizontal lines indicate the group median. Boxes represent the interquartile ranges and whiskers span the minimum to maximum values. C. Neutralizing antibody titers against RSV and hMPV. Graphs show half-maximal inhibitory dilutions (ID50). Horizontal lines indicate the group median. Boxes represent interquartile ranges and whiskers span the minimum to maximum values. D. Relationship between preF-binding antibody levels and neutralizing titers for RSV and hMPV, analyzed using a non-parametric Spearman correlation. E. Relationship between RSV and hMPV neutralizing titers. A subset of 6 samples with moderate-to-high RSV and hMPV neutralizing titers (highlighted in blue) were selected for electron microscopy-based polyclonal epitope mapping (EMPEM). F. Summary heatmap of RSV and hMPV preF and postF binding (ED50) and neutralization (ID50) titers across all donors. Values were normalized for each assay on a 0-100 scale and represented using color gradients. Donors selected for EMPEM analysis are highlighted in bold.

All 20 donors exhibited detectable levels of preF- and postF-binding IgG antibodies in an ELISA, with median ED50 values of 3,966 and 1,368 for RSV, and 832 and 409 for hMPV, respectively (**Fig. 1B** and **Fig. S2A**). Circulating antibodies also showed functional neutralizing capacity, with median ID50 titers of 288 for RSV and 1,190 for hMPV (**Fig. 1C** and **Fig. S2B**). These pneumovirus-specific antibody levels reflect prior natural exposure, since RSV vaccines had not been approved at the time of sample collection. RSV neutralization titers correlated moderately with preF-binding antibodies (Spearman r = 0.62, *p* = 0.0036) but not with postF-binding antibodies (Spearman r = 0.39, *p* = 0.0883) (**Fig. 1D** and **Fig. S2C**). In contrast, hMPV neutralization titers correlated strongly with both preF- and postF-binding antibodies (Spearman r = 0.86, *p* < 0.0001 and Spearman r = 0.78, *p* < 0.0001, respectively) (**Fig. 1D** and **Fig. S2C**). Consistent with these findings, hMPV preF- and postF-binding titers were also strongly associated with each other (Spearman r = 0.91, *p* < 0.0001) (**Fig. S2D**), which suggests that shared surface epitopes between the two conformations may be immunodominant. A subset of six donors exhibited moderately high preF-binding and neutralizing titers against both RSV and hMPV (**Fig. 1E-F**), making them suitable candidates for further structural characterization.

### Predicting immunodominance on pneumovirus preF proteins

At least seven antigenic sites have been identified on RSV and hMPV F proteins, including six analogous sites (Ø, I, II, III, IV and V) and one site specific to each virus (VI in RSV and the 66- 87 helix in hMPV)^12–14^. Sites I, II, III and IV, as well as the hMPV-specific 66-87 helix, are recognized in both the preF and postF conformations, whereas the remaining sites are uniquely displayed on preF (**Fig. 2A**). Understanding why some antigenic sites are targeted more frequently than others remains an important challenge in antibody immunology. To investigate whether intrinsic features of pneumovirus F proteins may contribute to a particular pattern, we applied the recently developed antigen surface immunodominance (ASI) model, which integrates the geometric, chemical and steric/spatial properties of antigens to predict immunodominant and subdominant regions on viral antigens^27^. We used the model to predict the relative immunodominance of the major antigenic sites on hMPV F and compared the results with those for RSV F^27^. This and subsequent analyses focused on the preF conformation, the main target of neutralizing antibodies.

**Figure 2.**
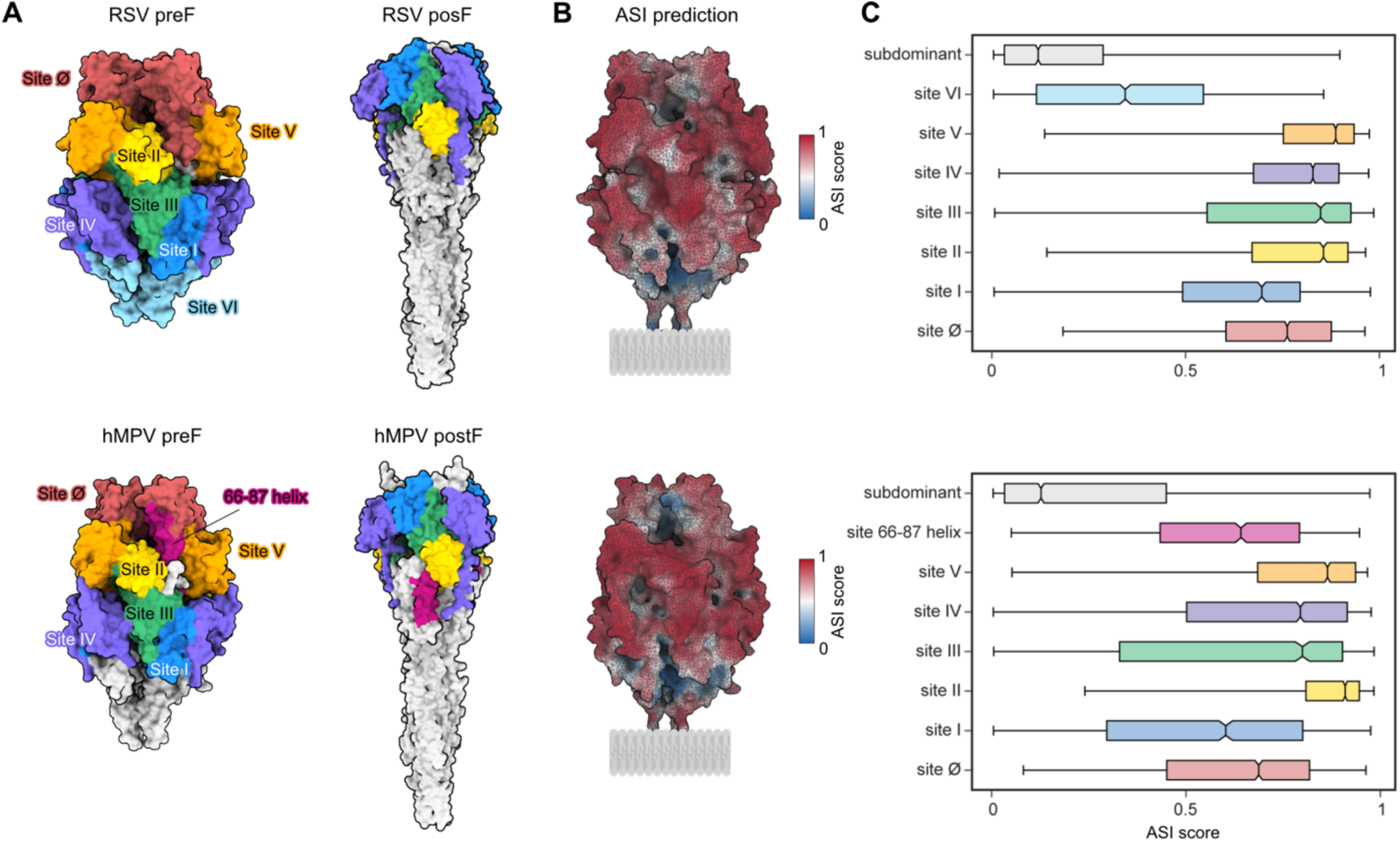
RSV and hMPV preF antigenic sites and immunodominance prediction using the antigen surface immunodominance (ASI) model. A. 3D models of RSV preF (SC-TM, PDB: 5C6B)^28^, RSV postF (PDB: 3RRR)^29^, hMPV preF (MPV-2c, PDB: 8VT2)^30^ and hMPV postF (PDB: 5L1X)^31^ colored according to previously defined antigenic sites^12–14^. B. ASI model prediction mapped onto the surface of the RSV and hMPV preF 3D models, simulated to be anchored on a viral membrane through a linker. **C.** Distribution of predicted ASI scores across the main antigenic sites in RSV and hMPV preF. The remaining amino acid residues were classified as ‘subdominant’. Boxes represent interquartile ranges and whiskers span the minimum to maximum values.

Overall, RSV and hMPV preF displayed highly similar ASI profiles. Large portions of the protein surface, including antigenic sites Ø, II, III, IV and V, were predicted to be highly immunogenic (**Fig. 2B-C**). In contrast, relatively low ASI scores were observed exclusively for surfaces not commonly targeted by mAbs (classified as ‘subdominant’), and in the case of RSV, for antigenic site VI. These lower scores likely reflect limited accessibility due to proximity to the viral membrane. Consistent with this interpretation, analysis of the individual features incorporated into the ASI model revealed that surface clearance (a measure of steric accessibility), followed by shape index (a descriptor of local surface curvature), most clearly distinguished immunodominant from subdominant regions (**Fig. S3A-B**).

Protrusion (a measure of surface exposure) was the feature most strongly associated with differences in predicted immunodominance among antigenic sites within each protein (**Fig. S3A- B**). Only subtle differences were observed between the two proteins, including a modestly lower ASI score for hMPV site Ø relative to RSV (**Fig. 2B-C**). This difference was predicted by the model to arise primarily from differences in proximity to potential N-linked glycosylation sites (PNGSs) (**Fig. S3A-B**). Based on the ASI predictions, most of the accessible antigenic surface in both preF proteins appears to feature optimal geometry coupled with low glycan shielding, resulting in a high probability of being targeted. However, these predictions are based solely on intrinsic antigen features and do not account for host-specific factors, such as infection history or immune status, that may shape antibody responses. We therefore re-examined published datasets and mapped the epitopes targeted by circulating human antibodies using site-specific competition assays and EMPEM.

### Pneumovirus-specific memory B cells and circulating antibodies exhibit broad preF specificities

Several high-throughput memory B cell profiling studies have characterized hundreds of RSV preF-specific mAbs^32–36^. These analyses revealed remarkably diverse antibody repertoires, with mAbs from humans and non-human primates recognizing all major antigenic sites (**Fig. 3A-B**). Notably, neutralization profiles of these mAbs allowed to establish a well-defined hierarchy of sensitivity, with antigenic sites Ø and V, located at the apex of the protein, being the most sensitive to antibody-mediated neutralization^32^ (**Fig. 3C**). Interestingly, the only study examining RSV mAbs from infants below three years of age showed a strong preference toward antigenic sites I and III, located in the basal and equatorial regions, respectively^33^. In contrast, adult-derived mAbs, particularly following vaccination with preF protein, seem to show a tendency to target antigenic sites Ø and V^35^.

**Figure 3.**
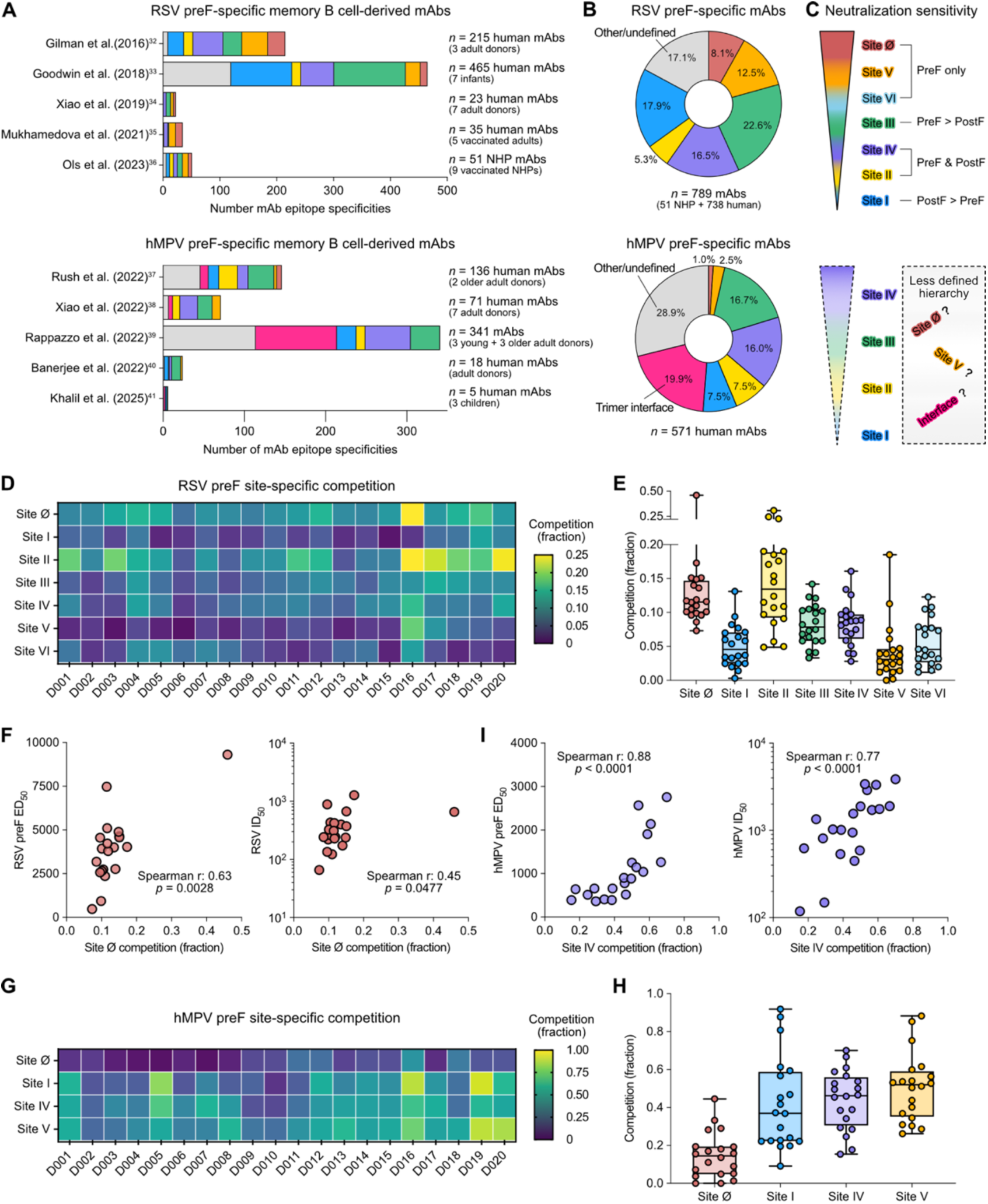
Pneumovirus-specific antibodies exhibit broad preF specificities. A. Specificities of RSV and hMPV mAbs cloned from preF-specific memory B cell receptors, based on data reported in previously published B cell profiling studies^32–41^. The number of specificities may exceed the number of mAbs in some studies, as the binding of some mAbs spans multiple antigenic sites, each of which is counted separately. **B.** Summary of RSV and hMPV mAbs specificities from all studies in **A** combined. **C.** RSV neutralization sensitivity hierarchy as established from preF-specific memory B cell profiling studies. Preferential recognition of the preF, postF or both conformations by antibodies targeting the respective sites is indicated. No definitive neutralization sensitivity hierarchy has yet been established for hMPV, but a tentative hierarchy is shown based on the most comprehensive hMPV study to date^39^. **D.** RSV preF site-specific targeting of circulating antibodies determined by a competition ELISA covering antigenic sites Ø, I, II, III, IV, V and VI. Competition levels higher than 0.25 were set to 0.25 for visualization purposes. **E.** Distribution of RSV preF-specific competition by circulating antibodies shown per antigenic site. Horizontal lines indicate the medians. Boxes represent interquartile ranges and whiskers span the minimum to maximum values. **F.** Relationship between levels of RSV preF site Ø competition with preF-binding titers and neutralizing titers, analyzed using a non-parametric Spearman correlation. **G.** hMPV preF site-specific targeting of circulating antibodies determined by a competition ELISA covering antigenic sites Ø, I, IV and V. **H.** Distribution of hMPV preF-specific competition by circulating antibodies shown per antigenic site. Horizontal lines indicate the medians. Boxes represent interquartile ranges and whiskers span the minimum to maximum values. **I.** Relationship between levels of hMPV preF site IV competition with preF-binding antibodies and neutralizing titers, analyzed using a non-parametric Spearman correlation.

Several hundreds of hMPV-specific mAbs have also been isolated from memory B cells. However, a definitive hierarchy of neutralization sensitivity has yet to be established, although available data suggests that sites III and IV may represent particularly vulnerable targets for neutralizing antibodies (**Fig. 3C**). Profiling of these mAbs also revealed broad diversity, with human antibodies recognizing all known sites, including the 66-87 helix located within the trimer interface^37–41^ (**Fig. 3A-B**). In contrast to RSV, most memory B cell-derived hMPV mAbs from adults preferentially bind sites located in the equatorial region rather than the apex. Remarkably, only a few hMPV- specific mAbs targeting site Ø or site V have been reported.

To determine which antigenic sites are most frequently targeted by RSV-specific circulating antibodies, we performed a competition ELISA using a panel of site-specific reference mAbs covering antigenic sites Ø, I, II, III, IV, V and VI (D25^42^, ADI-14457^32^, Palivizumab^29^, MPE8^43^, ADI- 18992^32^, hRSV90^44^ and 2.4K^14^, respectively) (**Fig. S4A**). The competition assay revealed that, similar to isolated mAbs, the circulating antibody repertoire of adults is diverse and targets all known antigenic sites (**Fig. 3D**). Although the overall level of competition varied across donors (**Fig. S4B**), sites Ø and II were among the most frequently targeted epitopes (**Fig. 3E**). Site Ø competition showed a positive correlation with both preF-binding titers and RSV neutralization (Spearman r = 0.63, *p* = 0.0028 and Spearman r = 0.45, *p* = 0.0477, respectively) (**Fig. 3F**), in line with the known importance of this preF-exclusive epitope as a major determinant of neutralizing activity. Importantly, site Ø competition did not correlate with postF-binding titers (**Fig. S4C-D**), further supporting that this association is driven by preF-specific antibody responses rather than overall F-reactive antibody levels.

We then characterized the antigenic site specificity of circulating hMPV antibodies using a similar setup, with ADI-61026^39^, DS7^45^, MPV482^40^, MPE8^43^, ADI-18992^32^, SAN27-14^37^ and MPV458^46^ as reference mAbs for hMPV preF sites Ø, I, II, III, IV, V and the 66-87 helix, respectively (**Fig. S4A**). Sites II, III and the 66-87 helix were excluded from further analysis, as self-competition could not be measured reliably with the corresponding unconjugated mAbs. Similar to RSV, the circulating antibody repertoire against hMPV preF was broad and recognized all antigenic sites that could be assessed (**Fig. 3G** and **Fig. S4B**). However, in contrast to RSV, site Ø was generally weakly targeted. Instead, stronger antibody responses were directed toward antigenic sites I, IV and V (**Fig. 3H**). Among these, site IV competition showed the strongest positive correlation with preF- and postF-binding titers (Spearman r = 0.88, *p* < 0.0001 and Spearman r = 0.89, *p* < 0.0001, respectively), as well as with hMPV neutralization (Spearman r = 0.77, *p* < 0.0001) (**Fig. 3I** and **Fig. S4E-F**), suggesting that antibodies targeting epitopes shared between the preF and postF conformations make an important contribution to hMPV neutralization. Our experimental findings revealed that circulating antibody repertoires against both RSV and hMPV preF are diverse and target multiple antigenic sites, rather than being strongly focused on a single epitope. This pattern is thus generally consistent with the ASI model predictions.

### EMPEM analysis of RSV preF-fragment antigen-binding (Fab) complexes reveals recurrent detection of apex-targeting antibodies

The subset of six donors with moderately high neutralizing titers against both pneumoviruses was further analyzed using EMPEM to structurally characterize their circulating antibody response. Full-length polyclonal IgGs were purified from plasma and enzymatically digested into Fabs that were then used to assemble the immune complexes (**Fig. S5A-B**). When polyclonal Fabs were complexed with RSV preF, only minor shifts to the left were observed in size-exclusion chromatography (SEC) traces compared to unbound preF, indicating an overall low abundance of RSV-specific antibodies (**Fig. S5C**). Although mentioned here in the context of the RSV dataset, this pattern was also observed for hMPV preF (**Fig. S5D**). Negative-stain EM image analysis further confirmed the low Fab occupancy level, showing that most particles corresponded to unbound preF, followed by complexes displaying one Fab bound and only occasionally complexes with two or three Fabs bound (**Fig. S6A-B**). Nevertheless, these data enabled the analysis of the most prevalent classes of polyclonal antibodies targeting the preF proteins.

The specific antigenic sites targeted by polyclonal Fabs were determined after visual inspection of RSV preF models colored by antigenic site and fitting them into the preF-Fab 3D reconstructions (**Fig. S7**). In all six donors, antibodies targeting site Ø at the apex of RSV preF were detected. These antibodies exhibited diverse angles of approach to the same epitope. Additional specificities were observed in two donors (D003 and D016), with antibodies recognizing the equatorial and upper middle regions (sites IV/V). Furthermore, antibodies against sites II/III or II/V were each identified in one donor (D001 and D004, respectively) (**Fig. 4A-C**). No antibodies were detected against sites I or VI located at the bottom of the protein (membrane-proximal). Although no clear correlation was observed between RSV neutralizing titers and epitope specificity, the consistent detection of site Ø antibodies across all donors is in line with the competition assay and suggests that the RSV neutralizing activity of circulating antibodies in adults is largely driven by this antibody class. Interestingly, EMPEM revealed that some site Ø- directed polyclonal antibodies, including those detected in donor D016, bound the apex epitope with an angle of approach resembling that of the RSV public antibody class represented by mAb 32.4K^35^ (**Fig. S8A**). Similarly, the site II/III-directed antibodies identified in donor D001 exhibited a binding mode reminiscent of that observed for public MPE8-like antibodies^47^ (**Fig. S8B**).

**Figure 4.**
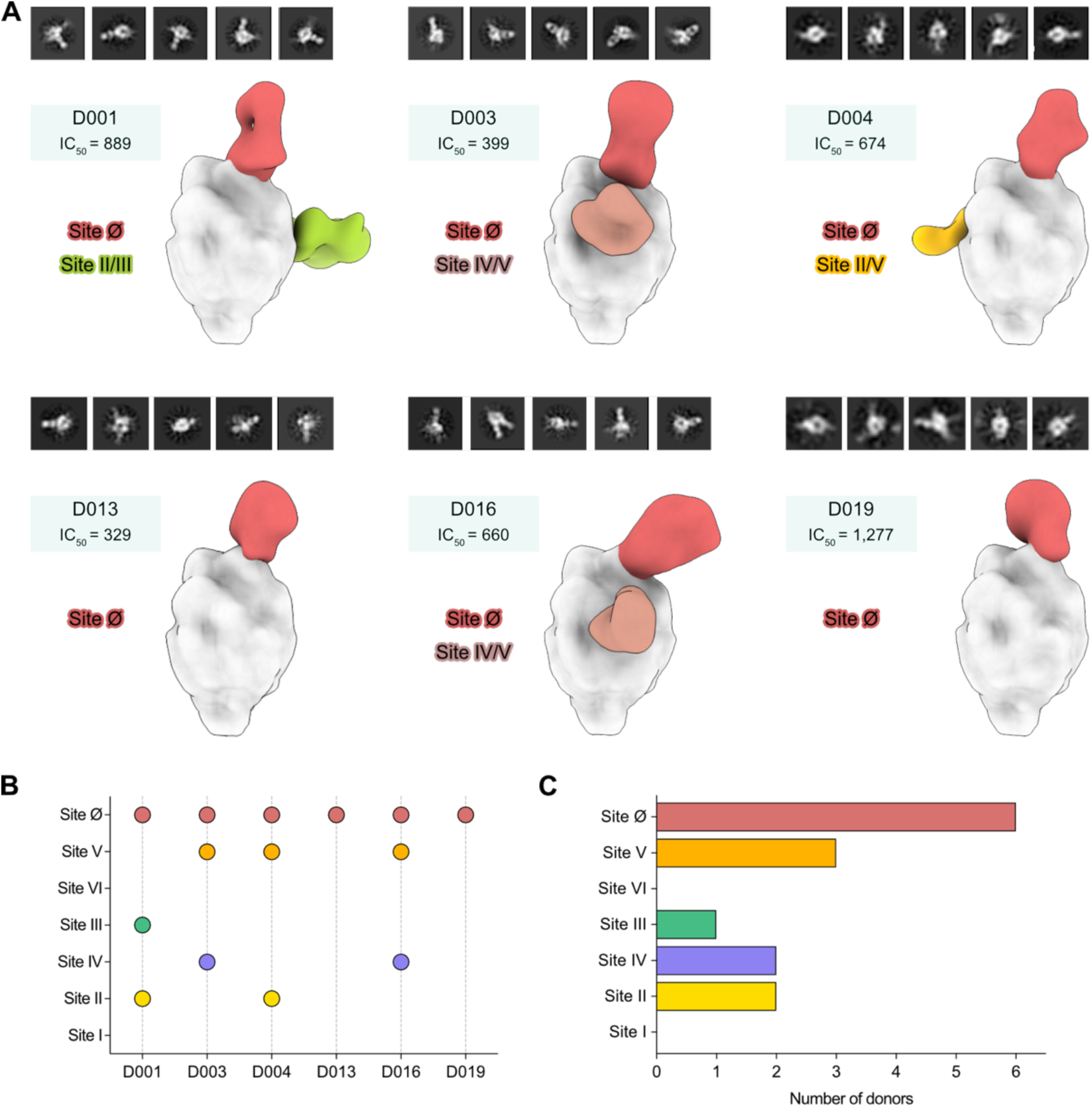
EMPEM analyses of antibody responses to RSV preF in healthy adult donors. A. Polyclonal antibody responses to RSV preF determined by negative-stain EMPEM in plasma from six donors with moderate RSV and hMPV neutralizing titers. Negative-stain EM 3D reconstructions of purified polyclonal Fabs bound to RSV preF. Representative 2D class averages are shown on top of each 3D reconstruction. The RSV neutralizing titers and antigenic sites targeted are also shown. **B.** Distribution of RSV preF antigenic sites targeted by the polyclonal antibody response per donor. **C.** Frequency of RSV preF antigenic site targeting by the polyclonal antibody response, summarized per antigenic site.

### EMPEM analysis of antibody responses to hMPV preF

Similarly to RSV, site Ø-specific antibodies targeting the apex of preF in diverse orientations were the most frequently detected class against hMPV preF, identified in four out of six donors. Among these four donors, two (D001 and D019) also had antibodies targeting sites II/V. Donor 001 additionally exhibited antibodies specifically targeting site V, resulting in the antibody response with highest observable epitope diversity within the cohort (**Fig. 5A-C**). Notably, antibodies from donor D001 also displayed the strongest neutralization capacity against hMPV. However, clear correlations between epitope specificity and neutralizing titers remain difficult to establish due to the relatively low number of antibodies recovered by EMPEM. For example, donors D013 and D016 had no reconstructable Fab densities despite exhibiting comparable or slightly higher neutralizing titers than donors D003, D004 and D019, for whom antibodies could be reconstructed. Although 3D reconstructions were not possible for D013 and D016, multiple 2D class averages provided evidence of polyclonal Fabs bound to hMPV preF, but in insufficient numbers to generate reliable 3D models (**Fig. 5A**).

**Figure 5.**
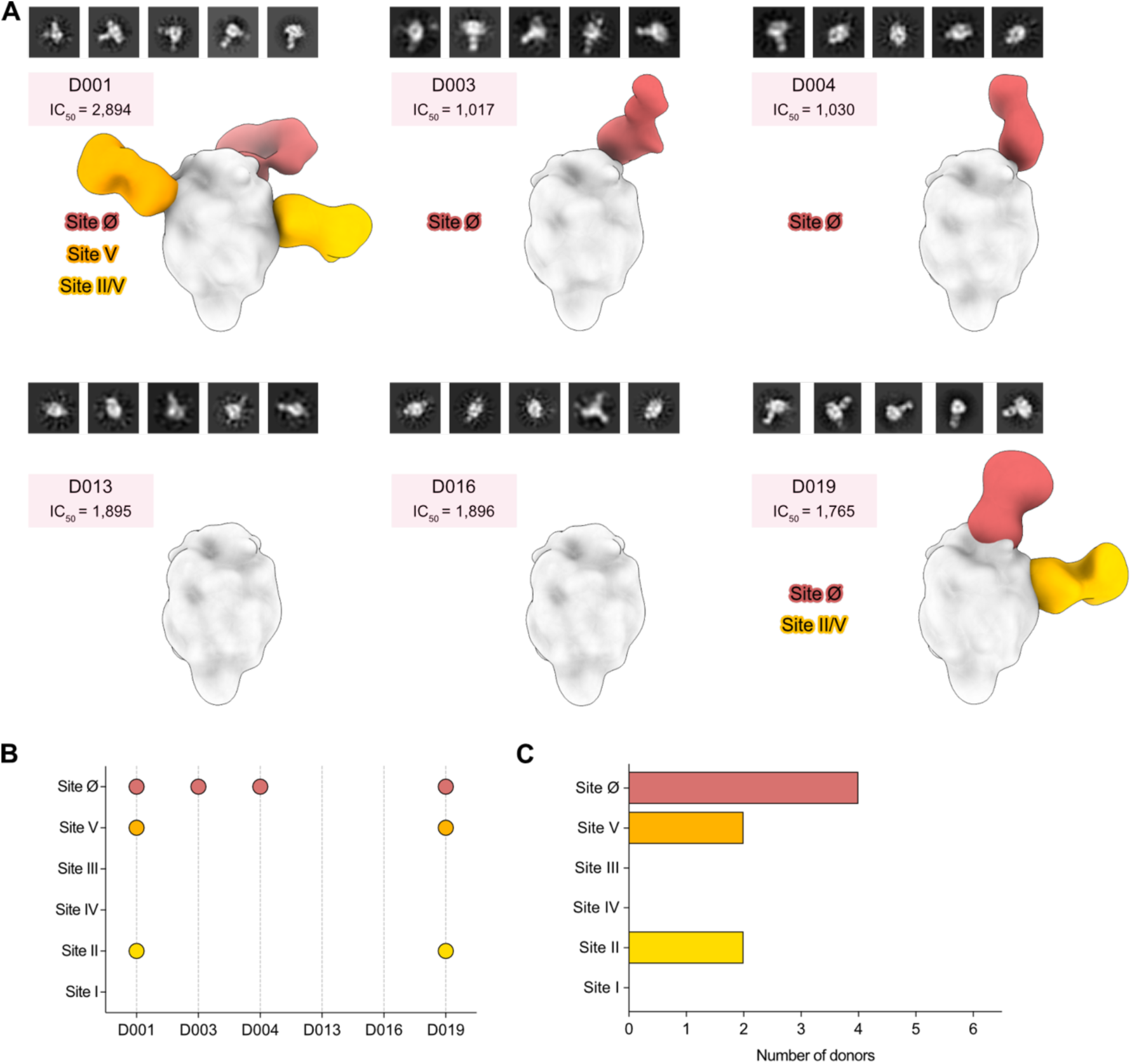
EMPEM analyses of antibody responses to hMPV preF in healthy adult donors. A. Polyclonal antibody responses to hMPV preF determined by negative-stain EMPEM in plasma from six donors with moderate RSV and hMPV neutralizing titers. Negative-stain EM 3D reconstructions of purified polyclonal Fabs bound to hMPV preF. Representative 2D class averages are shown on top of each 3D reconstruction. The hMPV neutralizing titers and antigenic sites targeted are also shown. **B.** Distribution of hMPV preF antigenic sites targeted by the polyclonal antibody response per donor. **C.** Frequency of hMPV preF antigenic site targeting by the polyclonal antibody response, summarized per antigenic site.

### Antibody-mediated pneumovirus preF destabilization

Among the different 3D reconstructions in our datasets, we occasionally encountered maps that showed a well-defined representation of the antigen, except for the apex-proximal region, where small protruding structures were present, often with one of those structures extending considerably (**Fig. S9A-B**). Although the resolution achieved by negative-stain EM does not allow definitive identification of these features, we observed that the extended protruding arm may resemble a Fab density and that its position and orientation are inconsistent with an intact, closed trimer. Based on these findings, we hypothesize that these maps represent antibody-mediated preF destabilization, where antibody binding to the closed trimer triggers partial dissociation into an open destabilized state that may affect the structural transitions necessary for membrane fusion, potentially contributing to neutralization. In addition, we observed structures resembling Fabs bound to preF in a monomeric state (**Fig. S10A**-**B**). Whether these monomeric forms arise from the intrinsic instability of preF or result from complete destabilization upon antibody binding cannot be determined from our data, but we mention them here for completeness.

## DISCUSSION

Here, we characterize the baseline polyclonal antibody response against RSV and hMPV in healthy donors. Consistent with previous studies^48–50^, we found that RSV preF-specific antibody levels correlate more strongly with virus neutralization than postF-specific antibodies. In contrast, for hMPV, both preF- and postF-specific antibody levels showed strong correlations with neutralizing activity, suggesting that antibodies recognizing both conformations contribute substantially to virus neutralization. Both competition ELISA and EMPEM analyses indicate that circulating antibodies target multiple antigenic sites on preF of both viruses, reflecting a broadly distributed polyclonal response upon repeated natural exposure. This pattern aligns well with ASI model predictions that several regions of the preF surface are important targets of antibody recognition and collectively shape its overall antigenic landscape.

Within the polyclonal repertoire, EMPEM consistently revealed antibodies binding to site Ø at the apex of preF for both RSV and hMPV, suggesting the presence of high-affinity antibodies to this neutralization-sensitive epitope across donors. For RSV, the predominant targeting of the preF apex is consistent with the specificities commonly observed among mAbs isolated from RSV- specific circulating memory B cells in adults^32,34,35^. By comparison, site Ø-directed responses to hMPV preF appear less common in both our competition ELISA and published memory B cell analyses^37–41^. Nevertheless, the detection of site Ø antibodies in multiple donors demonstrates that these antibodies remain a detectable component of the circulating hMPV antibody repertoire despite their comparatively lower abundance. One possible explanation for this discrepancy is that site Ø-directed responses may be less clonally diverse or exhibit more convergent modes of recognition, facilitating their reconstruction by EMPEM.

The low frequency of hMPV-specific apex-targeting antibodies has traditionally been attributed to the presence of two glycans (Asn57 and Asn172) shielding the apex of each F-protein protomer, which likely restrict antibody access to site Ø^51^. However, site Ø-specific hMPV antibodies able to penetrate the glycan shield have been isolated^37,39^, including one notable example (mAb 4F11) shown to approach the apex vertically and bind in a 1:1 Fab to trimer stoichiometry rather than the more typical 3:1 arrangement. This unique binding mode allows the antibody to insert between all apex-associated glycans^52^. Our EMPEM data supports the presence of apex-targeting hMPV-specific antibody classes within the circulating polyclonal repertoire and suggests that they may play an important role in hMPV neutralization, analogous to apex-directed responses observed for RSV.

Interestingly, our data revealed antibody-bound preF trimers that appeared partially or fully destabilized, providing evidence that the preF trimer is structurally dynamic (“breathes”) and can adopt conformations beyond the closed state. Antibody-mediated destabilization of the preF conformation has been documented previously for pneumoviruses, including examples involving site III-^53^ and site V-specific mAbs^54,55^. Similar antibody-driven effects have also been described for the measles virus preF protein^56^, a homologous class I fusion protein. In some of these cases, antibody binding increases trimer “breathing”, traps intermediate conformations or even triggers conversion to the postF state, thereby blocking the fusion machinery and contributing to neutralization. In the context of apex-targeting antibodies, binding may stabilize an open preF conformation and arrest fusion progression by restricting conformational changes in the refolding region 1. Whether the destabilized species observed here represent an example of this mechanism remains to be determined.

In addition to apex-targeting antibodies, our EMPEM analysis detected antibodies recognizing antigenic sites in the equatorial and upper middle region of preF (sites II, III, IV and V) in some donors. No antibodies directed to antigenic sites located at the base of the trimer (sites I or VI) were detected. Rather than reflecting a genuinely narrow antibody repertoire, this apparent limited diversity is more plausibly explained by inherent methodological constraints of the approach. EMPEM preferentially detects the most abundant and highest affinity antibody classes, making it less suited for capturing rare or lower affinity specificities. Consequently, the absence of base-targeting antibodies in our datasets should not be interpreted as evidence of their true absence in blood. These specificities were detected by competition ELISA but not by EMPEM, suggesting that they are present but fall below the detection threshold of EMPEM due to their lower affinity and/or more limited representation within the polyclonal repertoire. Future experiments incorporating photo-cross-linking of immune complexes to stabilize interactions with fast off-rate antibodies may help improve detection sensitivity^57^ and expand the range of pneumovirus antibody specificities resolvable by EMPEM.

Notably, the binding modes of some polyclonal antibodies resembled those of previously described public antibody classes, including the classes represented by mAbs 32.4K (site Ø)^35^ and MPE8 (site III)^47^. Within the EMPEM dataset, although site V-directed responses were observed, they did not exhibit features consistent with the public hRSV90-like antibody class^44^ and no site VI-directed (2.4K-like)^14^ responses were detected. Nevertheless, the identification of two known public antibody responses further supports the broader relevance of the antibody landscape observed here. While our goal in the present study was to characterize circulating antibody specificities in adults naturally exposed to pneumoviruses, future studies will be important to determine how pre-existing antibody repertoires in adults are reshaped by vaccination with pneumovirus preF vaccines. Such work could reveal whether vaccination preferentially expands antibodies targeting particular preF epitopes, especially those associated with high neutralization sensitivity, or instead promotes the overall diversity of the circulating antibody repertoire. Through this study, we have established a set of methods and a baseline map of epitope targeting within naturally acquired pneumovirus antibody responses to facilitate these future comparisons.

## Limitations of the study

Our study was limited by the relatively small number of donors for whom we could perform detailed EMPEM analysis, however, this is within the range of sample sizes examined in prior studies using this approach^20,21,23,58^. Importantly, the recurrent detection of similar antibody specificities across most individuals suggests that the observed responses are broadly representative of those likely present in larger populations. Another important consideration is that pneumoviruses display a heterogeneous mixture of F protein conformations on their surface, including trimeric preF and postF forms, as well as preF monomers (open trimers)^54^. Consequently, the antibody repertoire elicited during natural infection reflects exposure to multiple antigenic forms of F rather than preF trimers alone. This complicates the interpretation of epitope prevalence and apparent immunodominance. For example, antibodies targeting the equatorial or lower middle region may not arise solely because these sites are readily accessible on preF, but also because these epitopes are shared between preF and postF conformations. Furthermore, antibodies directed against such antigenic sites may preferentially recognize postF, potentially leading to their underrepresentation in analyses performed with preF antigens, particularly in structural approaches such as EMPEM.

## METHODS

### Cell lines

HEp-2 cells (ATCC: CCL-23; human carcinoma cell line) and LLC-MK2 cells (ATCC: CCL-7; rhesus monkey kidney cell line) were grown at 37°C and 5% CO2 in minimum essential media (MEM) supplemented with 10% fetal calf serum (FCS) and 1% penicillin/streptomycin. Expi293F cells (Gibco, CVCL_D615; human kidney cell line) were grown in suspension at 37°C, 8% CO2, 70-80% relative humidity and 100-125 rpm orbital shaking, in Expi293 expression medium supplemented with 1% penicillin/streptomycin.

### Viruses

Recombinant human RSV A2 strain line19F expressing the red fluorescent protein mKate2^59^ was propagated in HEp-2 cells from plaque purified stocks and harvested 4 days post-infection. Recombinant hMPV A2 strain CAN97-83 expressing green fluorescent protein (GFP)^60^ was propagated in LLC-MK2 cells from working stocks and harvested 4-5 days post-infection.

### Expression and purification of pneumovirus F proteins

Codon-optimized sequences for RSV preF (SC-TM)^28^, RSV postF^29^, hMPV preF (MPV-2c)^30^ and hMPV postF^31^ were synthesized and cloned into CMVR_HA1 plasmids by GenScript (Rijswijk, The Netherlands). Sequences for RSV preF and hMPV preF were designed to include Twin-Strep and Avi tags at the C-terminus. The RSV postF construct instead contains 8x His and Avi tags, while the hMPV postF construct contains all three Twin-Strep, Avi and 8x His tags. Recombinant proteins were expressed in Expi293F cells (viable cell density of 3 x 10^6^ cells/mL) upon transient transfection with 1 µg plasmid DNA/mL of culture using either the ExpiFectamine 293 transfection kit (Gibco) for RSV preF, hMPV preF and hMPV postF or 3 µg/mL of PEI MAX MW 40,000 (Polysciences) for RSV postF. Except for RSV preF, all other constructs were co-transfected with a furin plasmid in a 1:4 furin to target plasmid ratio. Culture supernatants were harvested 4-5 days post-transfection, clarified by centrifugation and filtered through a 0.22 µm vacuum-driven sterile filter. The supernatant of RSV postF was pH- and salt-stabilized to a final concentration of 10 mM PBS pH 7.4, while supernatants of Twin-Strep-tagged proteins were stabilized to a final concentration of 100 mM Tris/HCl pH 8.0, 150 mM NaCl, 1 mM EDTA. Finally, biotin present in Twin-Strep-tagged protein supernatants was blocked by incubation with BioLock (IBA Lifesciences) for 20 min at room temperature.

Proteins were purified using a two-step procedure consisting of affinity-based capture followed by SEC. RSV postF was captured by incubation with a Ni Sepharose excel resin (Cytiva) for 1 h at 4°C under head-to-heel rotation and eluted by gravity flow using 500 mM imidazole in PBS. Twin-Strep-tagged proteins were captured in-column with a Strep-Tactin Sepharose or a Strep- Tactin 4Flow high capacity resin (IBA Lifesciences) and eluted using 100 mM Tris/HCl pH 8.0, 150 mM NaCl, 1 mM EDTA, 2.5-10 mM desthiobiotin. Eluates were concentrated using Amicon Ultra centrifugal filters (Millipore) and further purified on a Superdex 200 Increase 10/300 GL column (Cytiva) equilibrated with either 100 mM Tris/HCl pH 8.0, 150 mM NaCl, 1 mM EDTA or 10 mM PBS pH 7.4. Purified preparations were spiked with 5% v/v final concentration of glycerol before flash-freezing.

Protein purity was assessed by standard SDS-PAGE and Coomassie staining under non-reducing conditions. Antigenic profiles and structural integrity were evaluated using ELISA and negative-stain EM, respectively, as described below. Of note, the hMPV postF protein was heated at 70°C for 30 min before use to ensure complete conversion to the postF conformation.

### Negative-stain EM sample preparation and data collection (antigen characterization)

Samples were diluted to 0.01 mg/mL from starting concentrations in 25 mM Tris pH 7.5, 100 mM NaCl. 3 µL of diluted sample was applied to a grid and given 30 s to adsorb. The protein sample was then wicked using filter paper. Samples were then stained with 2% w/v uranyl formate and given 30 s to adsorb. The stain was then wicked with the filter paper. This staining process was repeated three times with residual moisture air drying following final application of filter paper. Proteins were imaged using Talos L120C TEM instruments (Thermo Scientific). For each sample, approximately 150–200 micrographs were collected at a total electron dose of 50 e⁻/Å².

### Negative-stain EM data processing and analysis (antigen characterization)

Data processing was performed using CryoSPARC^61^. CTF parameters were estimated using Patch CTF Estimation with default settings. Initial particle picking was performed using Blob Picker with a particle diameter range of 150–270 Å, followed by manual inspection of particle picks. Particles were extracted using a box size of 160 pixels and subjected to 2D classification into 50 classes. Representative 2D classes were selected as templates for a second round of particle picking using a particle diameter of 270 Å. Particle picks were manually inspected and filtered to retain high-quality particles. Selected particles were subjected to *ab initio* 3D reconstruction with C3 symmetry imposed and three output classes specified. The class most closely resembling trimeric RSV preF or hMPV preF was selected for subsequent non-uniform refinement, yielding final reconstructions at resolutions of approximately 15–20 Å. Reference structures of RSV preF (PDB ID: 5C6B)^28^ or hMPV preF (PDB ID: 8VT2)^30^ were rigid-body docked into their corresponding reconstructed maps for structural analysis. Maps were flipped along the Z-axis as needed to establish the correct handedness relative to the reference structures.

### Human samples

Blood samples from anonymous healthy adult donors were obtained from the Karolinska University Hospital, Stockholm, Sweden. Donors were considered healthy according to standard blood donation eligibility criteria, which typically includes being 18-65 years old, weighing ≥50 kg, having no active illness or infection and having no recent high-risk exposures. No additional demographic or clinical information was available. Plasma was isolated using standard centrifugation-based separation techniques and heat-inactivated at 56°C for 1 h before its use. The study was conducted within the scope of the ethics permit No. 2025-01799-02, issued by the Swedish Ethical Review Authority.

### ELISA (antigenicity of pneumovirus F proteins)

Antigenic profiles of RSV preF (SC-TM)^28^, hMPV preF (MPV-2c)^30^ and hMPV postF^31^ were evaluated by ELISA using 96-well Strep-Tactin XT plates (IBA Lifesciences). Plates were first blocked with 200 µL/well of blocking buffer (2% bovine serum albumin [BSA] in 50 mM Tris/HCl, pH 7.4, 140 mM NaCl, supplemented with 0.05% v/v Tween 20 [TBS-T]) for at least 1 h. Then, 100 µL/well of Twin-Strep-tagged pneumovirus F antigens (1 µg/mL in TBS) was added and incubated for 1 h. Antigens were subsequently incubated for 2 h with 100 µL/well of five-fold serial dilutions (starting at 10 µg/mL) of a reference panel of RSV or hMPV mAbs prepared in blocking buffer. Conformation-specific mAbs were detected after 1 h incubation with 100 µL/well of HRP-conjugated goat anti-human IgG (Jackson ImmunoResearch) diluted 1:10,000-1:20,000 in blocking buffer. Plates were washed three times with washing buffer (TBS-T) between every step using an automatic plate washer. All incubations were performed at room temperature. Signal was developed upon addition of 1-Step Ultra TMB-ELISA substrate (Thermo Fisher Scientific).

The reaction was stopped after 3-5 min by addition of 1 M H2SO4. Absorbance was measured at 450 nm with background correction at 570 nm on a Varioskan LUX Multimode Microplate Reader (Thermo Scientific). Samples were analyzed in duplicate. Antibodies recognizing an alternative conformation or specific to the other virus were used as negative controls.

The antigenic profile of RSV postF^29^ was evaluated using conventional half-area 96-well plates directly coated with 50 µL/well of antigen (2 µg/mL in PBS) and incubated overnight at 4°C. Plates were then blocked with 100 µL/well of blocking buffer (5% w/v skim milk in PBS) for 1 h. The antigen was subsequently incubated for 2 h with 50 µL/well of 6-fold serial dilutions (starting at 20 µg/mL) of a reference panel of RSV mAbs prepared in blocking buffer. Conformation-specific mAbs were detected after 1 h incubation with 50 µL/well of HRP-conjugated goat anti-human IgG (Jackson ImmunoResearch) diluted 1:5,000 in blocking buffer. Plates were washed three times with washing buffer (PBS supplemented with 0.05% v/v Tween 20 [PBS-T]) between every step using an automatic plate washer. Except for the coating step incubated at 4°C, all other incubations were performed at room temperature. Signal development and absorbance measurements were performed as described above. Samples were analyzed in duplicate. An antibody recognizing an alternative conformation was used as a negative control.

### ELISA (circulating pneumovirus antibodies)

Circulating antibodies against RSV and hMPV F proteins were quantified by ELISA. For RSV preF and postF, half-area 96-well plates were coated with 50 µL/well of antigen (2 µg/mL in PBS) and incubated overnight at 4°C. Plates were then blocked with 100 µL/well of blocking buffer (5% w/v skim milk in PBS) for 1 h. Antigens were subsequently incubated for 2 h with 50 µL/well of four- fold serial dilutions of human plasma (starting at 1:25) prepared in blocking buffer. Conformation-specific antibodies were detected after 1 h incubation with 50 µL/well of HRP- conjugated goat anti-human IgG (Jackson ImmunoResearch) diluted 1:5,000 in blocking buffer. Plates were washed three times with washing buffer (PBS-T) between every step using an automatic plate washer. Except for the coating step incubated at 4°C, all other incubations were performed at room temperature. Signal development and absorbance measurements were performed as described above. Samples were analyzed in duplicate. Antibody titers are reported as the half-maximal effective dilution (ED50), determined using a four-parameter non-linear regression curve fit in Prism version 11 (GraphPad Software).

For hMPV preF and postF, half-area 96-well plates were coated with 50 µL/well of Strep-Tactin (2 µg/mL in PBS) and incubated overnight at 4°C. Plates were then blocked with 100 µL/well of blocking buffer (2% w/v BSA in PBS-T) for 1 h. Next, 50 µL/well of antigen (2 µg/mL in PBS) was added and incubated for 1 h. Following antigen immobilization, plates were subsequently incubated for 2 h with 50 µL/well of four-fold serial dilutions of human plasma (starting at 1:25) prepared in blocking buffer. Conformation-specific antibodies were detected after 1 h incubation with 50 µL/well of HRP-conjugated goat anti-human IgG (Jackson ImmunoResearch) diluted 1:15,000 in blocking buffer. Plates were washed three times with washing buffer (PBS-T) between every step using an automatic plate washer. Except for the coating step incubated at 4°C, all other incubations were performed at room temperature. Signal development and absorbance measurements were performed as described above. Samples were analyzed in duplicate. Antibody titers are reported as the half-maximal effective dilution (ED50), determined using a four- parameter non-linear regression curve fit in Prism version 11 (GraphPad Software).

### Competition ELISA

The epitope specificity of circulating antibodies against RSV and hMPV preF proteins was determined by a competition ELISA using a panel of site-specific reference mAbs. Reference mAbs for RSV preF included D25 (site Ø)^42^, ADI-14457 (site I)^32^, Palivizumab (site II)^29^, MPE8 (site III)^43^, ADI-18992 (site IV)^32^, hRSV90 (site V)^44^ and 2.4K (site VI)^14^. Reference mAbs for hMPV preF initially included ADI-61026 (site Ø)^39^, DS7 (site I)^45^, MPV482 (site II)^40^, MPE8 (site III)^43^, ADI-18992 (site IV)^32^, SAN27-14 (site V)^37^ and MPV458 (66-87 helix)^46^, although ultimately we could only reliably measure competition to sites Ø, I, IV and V.

For RSV preF, half-area 96-well plates were coated with 50 µL/well of antigen (2 µg/mL in PBS) and incubated overnight at 4°C. Plates were then blocked with 100 µL/well of blocking buffer (5% w/v skim milk in PBS) for 1 h. Next, plates were incubated with 25 µL/well of human plasma diluted 1:20 in blocking buffer for 30 min. Subsequently, 25 µL/well of biotinylated site-specific reference mAbs diluted in blocking buffer were added and incubated for 90 min. The concentration of each biotinylated mAb was predetermined from binding curves (4-fold serial dilution starting at 20 µg/mL) as the concentration yielding a background-subtracted signal of approximately 2-3 (based on a four-parameter non-linear regression curve fit). Site-specific antibodies were detected after 1 h incubation with 50 µL/well of Pierce High Sensitivity NeutrAvidin-HRP (Thermo Scientific) diluted 1:40,000 in PBS-T. Plates were washed three times with washing buffer (PBS-T) between each step using an automatic plate washer. Except for the coating step incubated at 4°C, all other incubations were performed at room temperature. Signal development and absorbance measurements were performed as described above. Samples were analyzed in duplicate. RSV reference mAbs were biotinylated using the EZ-Link Sulfo-NHS-LC-Biotin reagent (Thermo Scientific) following the manufacturer’s instructions.

For hMPV preF, half-area 96-well plates were coated with 50 µL/well of Strep-Tactin (2 µg/mL in PBS) and incubated overnight at 4°C. Plates were then blocked with 100 µL/well of blocking buffer (2% w/v BSA in PBS-T) for 1 h. Next, 50 µL/well of antigen (2 µg/mL in PBS) was added and incubated for 1 h. Following antigen immobilization, plates were incubated with 25 µL/well of human plasma diluted 1:20 in blocking buffer for 30 min. Subsequently, 25 µL/well of HRP- conjugated site-specific reference mAbs diluted in blocking buffer were added and incubated for 90 min. The concentration of each HRP-mAb was predetermined from binding curves (three-fold serial dilution starting at 20 µg/mL) as the concentration yielding a background-subtracted signal of approximately 2-3 (based on a four-parameter non-linear regression curve fit). Plates were washed three-six times with washing buffer (PBS-T) between each step using an automatic plate washer. Except for the Strep-Tactin coating step incubated at 4°C, all other incubations were performed at room temperature. Signal development and absorbance measurements were performed as described above. Samples were analyzed in duplicate. hMPV reference mAbs were conjugated to HRP using the HRP Conjugation Kit – Lightning-Link (abcam) following the manufacturer’s instructions.

In parallel to the analysis of human plasma, self-competition curves were generated for each biotinylated or HRP-conjugated reference mAb. Briefly, a three-fold serial dilution of the corresponding unconjugated mAb (starting at 10 µg/mL) was used to compete with binding of the biotinylated or HRP-conjugated mAb. The level of competition (reported as a fraction) was normalized by defining the signal obtained in the presence of 10 µg/mL of unconjugated mAb as 100% competition (fraction of 1) and the signal without unconjugated mAb as 0% competition (fraction of 0). Competition values of plasma samples for each antigenic site were then calculated by interpolation of the corresponding normalized self-competition curve. All curve fitting and interpolation were performed using a four-parameter non-linear regression in Prism version 11 (GraphPad Software).

### RSV neutralization assay

HEp-2 cells suspended in FluoroBrite Dulbecco’s Modified Eagle Medium (DMEM; Gibco) supplemented with 10% FCS were seeded in black 384-well optically clear bottom plates (Thermo Fisher Scientific) at a density of 6 x 10^3^ cells/well and incubated overnight at 37°C with 5% CO2. In standard 96-well cell culture plates, serial three-fold dilutions of plasma (starting at 1:10) prepared in culture medium were mixed with an equal volume of 1:4-8 diluted recombinant RSV A2 mKate^59^ and incubated for 1 h at 37°C. Subsequently, 50 µL of the plasma-virus mixture was added to pre-seeded HEp-2 cells and incubated for 26 h at 37°C with 5% CO2. Fluorescence (excitation at 588 nm and emission at 635 nm) was measured from the bottom on a Varioskan LUX Multimode Microplate Reader (Thermo Scientific). Data were normalized by defining maximum signal in wells without plasma as 100% infection and a fluorescence signal of 0.2 as 0% infection. Neutralizing titers were expressed as the half-maximal inhibitory dilution (ID50), calculated using a four-parameter non-linear regression curve fit in Prism version 11 (GraphPad Software).

### hMPV neutralization assay

LLC-MK2 cells suspended in FluoroBrite DMEM (Gibco) were seeded in black 384-well optically clear bottom plates (Thermo Fisher Scientific) at a density of 6 x 10^3^ cells/well and incubated overnight at 37°C with 5% CO2. In standard 96-well cell culture plates, serial three-fold dilutions of plasma (starting at 1:60) prepared in culture medium were mixed with an equal volume of 1:8 diluted recombinant hMPV A2 GFP^60^ and incubated for 1 h at 37°C. Subsequently, 50 µL of the plasma-virus mixture was added to pre-seeded LLC-MK2 cells and incubated for 48 h at 37°C with 5% CO2. Fluorescence (excitation at 489 nm and emission at 511 nm) was measured from the bottom on a Varioskan LUX Multimode Microplate Reader (Thermo Scientific). Data were normalized by defining maximum signal in wells without plasma as 100% infection and a fluorescence signal of 0.2 as 0% infection. Neutralizing titers were expressed as the half-maximal inhibitory dilution (ID50), calculated using a four-parameter non-linear regression curve fit in Prism version 11 (GraphPad Software).

### Plasma IgG purification

Plasma samples from individuals selected for EMPEM analysis were diluted 5-fold in PBS pH 7.4 and applied to CaptureSelect IgG-Fc (multispecies) affinity matrix (Thermo Scientific) equilibrated in the same buffer for in-column capture of polyclonal IgGs. The resin and undiluted plasma were used at a 1:1 ratio. After at least three washes with PBS pH 7.4, IgGs were eluted by gravity flow using 0.1 M glycine pH 2.5, immediately neutralized with 1 M Tris/HCl pH 8 and filtered through a 0.22-0.45 µm pore filter. Eluates were buffer exchanged into 100 mM Tris/HCl, 2 mM EDTA pH 8 using Amicon Ultra centrifugal filters (Millipore) with a 30 kDa cut-off.

### IgG digestion and Fab purification

IgGs were digested with freshly prepared papain from *Carica papaya* (Sigma-Aldrich) at 1 mg/mL in 100 mM Tris/HCl pH 8.0, 2 mM EDTA, 10 mM L-cysteine, activated by incubation at 37°C for 15 min. Digestion reactions proceeded for 4-5 h at 37°C, using 40 µL of activated papain per 1 mg of isolated IgG. Reactions were quenched with iodoacetamide at a final concentration of 30 mM, and digested samples were concentrated using Amicon Ultra centrifugal filters (Millipore) with a 10 kDa cut-off. Fab and Fc fragments were separated from undigested IgG and F(ab’)2 fragments by SEC on a Superdex 200 Increase 10/300 GL column (Cytiva) equilibrated with 100 mM Tris/HCl pH 8, 150 mM NaCl, 1 mM EDTA. Purified Fab and Fc fractions were concentrated using Amicon Ultra centrifugal filters (Millipore) with a 10 kDa cut-off. Purity of the fractions of interest was assessed by standard SDS-PAGE and Coomassie staining under non-reducing conditions.

### Pneumovirus preF-polyclonal Fab immune complexes

Immune complexes were prepared by mixing 10 µg of RSV preF or hMPV preF with 750 µg of purified polyclonal Fab/Fc. Mixtures were incubated overnight at room temperature. Complexes were subsequently purified from unbound Fab/Fc by SEC on a Superdex 200 Increase 10/300 GL column (Cytiva) equilibrated with 20 mM Tris/HCl pH 7.4, 150 mM NaCl. Purified complexes were concentrated using Amicon Ultra centrifugal filters (Millipore) with a 30 kDa cut-off and used immediately for preparing EM grids.

### Negative-stain EM sample preparation and data collection (EMPEM)

Purified immune complexes were diluted to approximately 20 µg/mL in 20 mM Tris/HCl pH 7.4, 150 mM NaCl and applied onto glow-discharged (3 mA for 30 s) carbon film 300-mesh copper grids (CF300-Cu-50, Electron Microscopy Sciences). Samples were allowed to adsorb for 15-60 s and stained with 2% w/v uranyl formate for 60 s. After each step, excess liquid was blotted with filter paper and the grids were ultimately let air dry. Stained grids were imaged on a Talos L120C electron microscope equipped with a Ceta camera (Thermo Scientific) operating at 120 keV and 57,000x nominal magnification (2.44 Å/pixel). Automated data acquisition was performed using EPU software (Thermo Scientific). For each dataset, between 100 and 300 micrographs were acquired.

### Negative-stain EM data processing and analysis (EMPEM)

All datasets were imported into CryoSPARC^61^ version 4.5.3 for pre-processing, including particle picking, inspection, extraction and 2D classification. For each sample, between 174k and 472k particles were extracted. After at least two rounds of 2D classification, selected 2D classes corresponding to preF-Fab complexes or preF alone were transferred to Relion^62^ version 4.0.1 for CTF estimation and additional 2D classification. Selected 2D classes, accounting for between 80k and 192k particles, were further processed through iterative rounds of 3D classification, where similar-looking classes were combined and reclassified. A low-resolution model of unbound pneumovirus preF was used as a reference for 3D steps. 3D classes with well-defined Fab reconstructions underwent 3D auto-refinement in Relion^62^ version 5.0.1. Fab volumes from representative 3D reconstructions of antigen-specific polyclonal Fabs were segmented using the *Segger* tool^63^ and incorporated into final composite 3D maps in UCSF ChimeraX^64,65^ version 1.12.

### Immunodominance predictions and ASI model scores

The ASI model, developed to predict immunodominance from geometric, chemical, steric and dynamic properties of protein surfaces, has been described previously^27^. In this study, the model was applied to an experimental structure of hMPV preF (MPV-2c; PDB 8VT2)^30^ and compared to a prediction for RSV preF^27^ (SC-TM; PDB 5C6B)^28^. A nanoparticle scaffold was linked to the bottom of the antigens to mimic their anchoring to a viral membrane. For each structure, a molecular surface representation was generated and ASI scores were computed for individual surface vertices. Vertex-level scores were subsequently aggregated and reported at the antigenic-site level. Antigenic sites on RSV preF were initially annotated based on previously published definitions^14^ and were subsequently refined to exclude buried residues. Antigenic sites on hMPV preF were annotated to correspond to the analogous sites on RSV preF, with the additional inclusion of the hMPV-specific 66-87 helix epitope^46^. Buried residues were also excluded.

### Data analysis and visualization

Prism version 11 (GraphPad Software) was used to generate plots and perform statistical analysis. Statistical tests are indicated in the corresponding figure legends. *p* values ≥ 0.05 were considered not significant. UCSF Chimera X^64,65^ version 1.12 was used for structure visualization and analysis.

### Data availability

Refined 3D reconstructions from the EMPEM analyses have been deposited in the Electron Microscopy Data Bank (EMDB; http://www.emdatabank.org) under accession codes EMD- 100673, EMD-100674, EMD-100675, EMD-100676, EMD-100679, EMD-100681, EMD-100682, EMD-100683, EMD-100684, EMD-100686, EMD-100689, EMD-100691, EMD-100692, EMD-100693, EMD-100694, EMD-100695 and EMD-100696 (**Table S1**) and will be released upon publication of the manuscript. The ASI prediction software will be deposited to https://github.com/UPANTANASIJEVIC/ASI.

## ACKNOWLEDGEMENTS

We gratefully acknowledge the contribution of the anonymous blood donors. We also thank all members from Leo Hanke’s (Karolinska Institutet) and Aleksandar Antanasijevic’s (École Polytechnique Fédérale de Lausanne) labs for generously sharing their lab space, equipment and expert technical help. We further thank Jolijn van der Geest and Mariana Prado Marmorato for their contributions during early stages of assay development, as well as André Nicolás León and Eric de Sousa for their expert technical support in setting up and configuring our data-processing workstation. This work was financially supported by the Swedish Research Council (Vetenskapsrådet; 2023-02396 and 2025-06582 to K.L.), the Swedish Society for Medical Research (SSMF; PG-25-0564-H-01 to E.B.M.) and the Swiss National Science Foundation (SNSF; grant 3200-0-239993 to A.A). E.B.M. gratefully acknowledges additional financial support by the European Federation of Immunological Societies (EFIS) and Immunology Letters for the short-term fellowship that made possible his visit to École Polytechnique Fédérale de Lausanne. The illustration in **Fig. 1A** was partially created with BioRender.com.

## AUTHOR CONTRIBUTIONS

Conceptualization: Erick Bermúdez-Méndez, Sebastian Ols, Marcos C. Miranda, Aleksandar Antanasijevic, Karin Loré.

Methodology: Erick Bermúdez-Méndez, Sebastian Ols, Shuhao Xiao, Florian Gegenfurtner, Marcos C. Miranda, Andrew J. Borst, Aleksandar Antanasijevic.

Investigation: Erick Bermúdez-Méndez, Sebastian Ols, Leonardo V. Cufaro, Shuhao Xiao, Zhitong Peng, Florian Gegenfurtner, Elise R. Eray, Marcos C. Miranda, Rodrigo Arcoverde Cerveira, Mikel Segura García, Miguel A. Véliz Pernilla, Aleksandar Antanasijevic.

Formal analysis: Erick Bermúdez-Méndez, Leonardo V. Cufaro, Shuhao Xiao, Zhitong Peng, Andrew J. Borst, Aleksandar Antanasijevic.

Visualization: Erick Bermúdez-Méndez, Shuhao Xiao. Resources: Leo Hanke, Neil P. King.

Supervision: Andrew J. Borst, Neil P. King, Aleksandar Antanasijevic, Karin Loré. Funding acquisition: Erick Bermúdez-Méndez, Aleksandar Antanasijevic, Karin Loré. Writing – original draft: Erick Bermúdez-Méndez

Writing – review & editing: all authors.

