## Supplementary Information for "Structural and functional characterization of polyclonal antibody responses reveals shared and distinct antigenic features of RSV and hMPV fusion proteins"

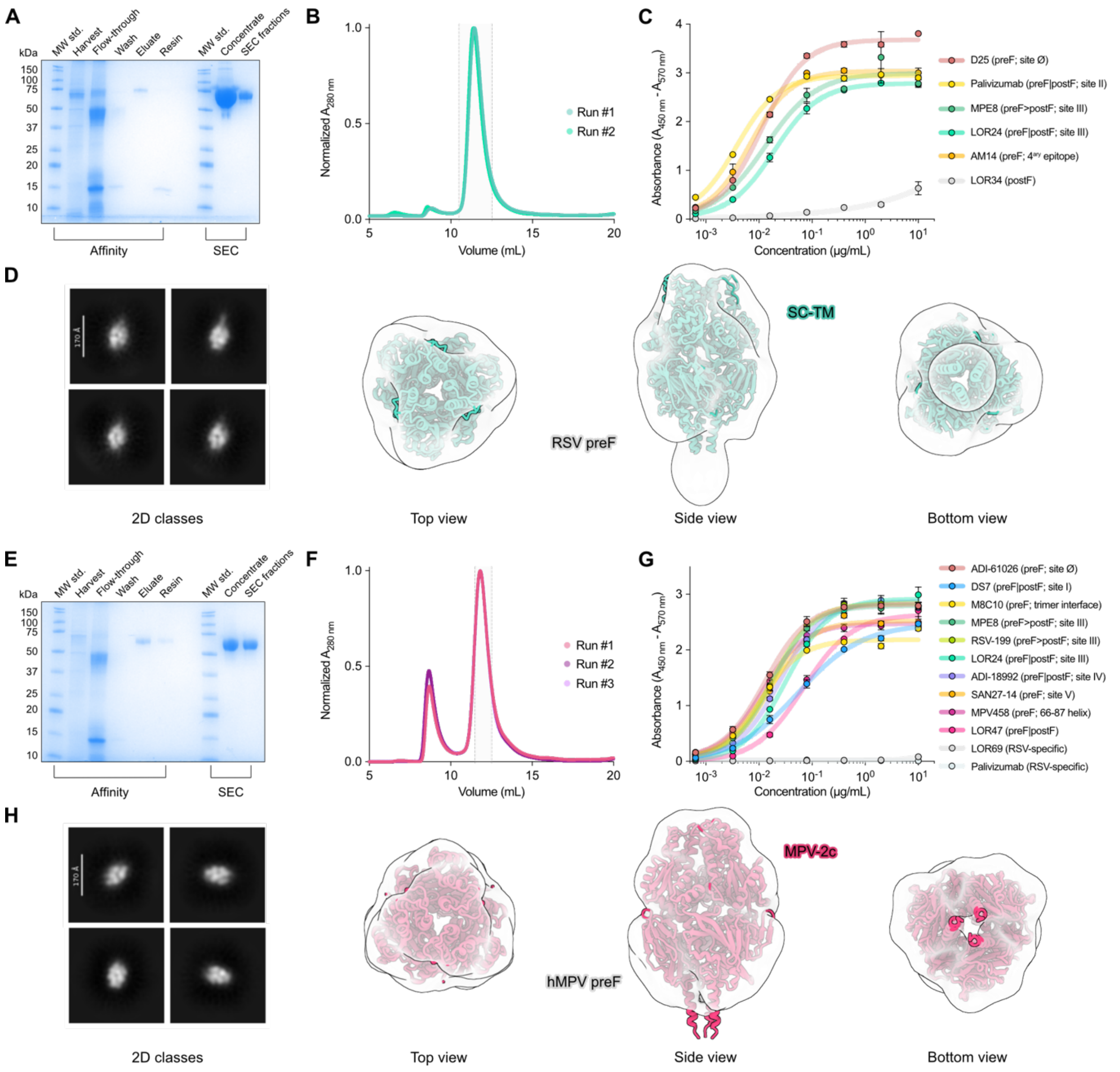


**Figure S1. Functional and structural validation of RSV and hMPV preF proteins.**

**A.** SDS-PAGE with Coomassie staining analysis of fractions collected during RSV preF (SC-TM, PDB: 5C6B)[1] purification by Strep-Tactin-based affinity chromatography (left) and size-exclusion chromatography (SEC; right).

**B.** SEC traces of Strep-Tactin-purified RSV preF on a Superdex S200 Increase 10/300 GL column. Peak of interest is highlighted in light gray. The overlay of two runs is shown.

**C.** Antigenicity profile of RSV preF assessed by ELISA using a panel of conformation-specific RSV reference monoclonal antibodies (mAbs). LOR34 is a postF-specific RSV mAb used as a negative control.

**D.** Structural validation of RSV preF analyzed by negative-stain EM. Representative 2D class averages and 3D maps with the structure of SC-TM fit into the map.

**E.** SDS-PAGE with Coomassie staining analysis of fractions collected during hMPV preF (MPV-2c, PDB: 8VT2)[2] purification by Strep-Tactin-based affinity chromatography (left) and SEC (right).

**F.** SEC trace of Strep-Tactin-purified hMPV preF on a Superdex S200 Increase 10/300 GL column. Peak of interest is highlighted in light grey. The overlay of three runs is shown.

**G.** Antigenicity profile of hMPV preF assessed by ELISA using a panel of conformation-specific hMPV reference mAbs. Palivizumab and LOR69[3] are RSV-specific mAbs used as negative controls.

**H.** Structural validation of hMPV preF analyzed by negative-stain EM. Representative 2D class averages and 3D maps with the structure of MPV-2c fit into the map.

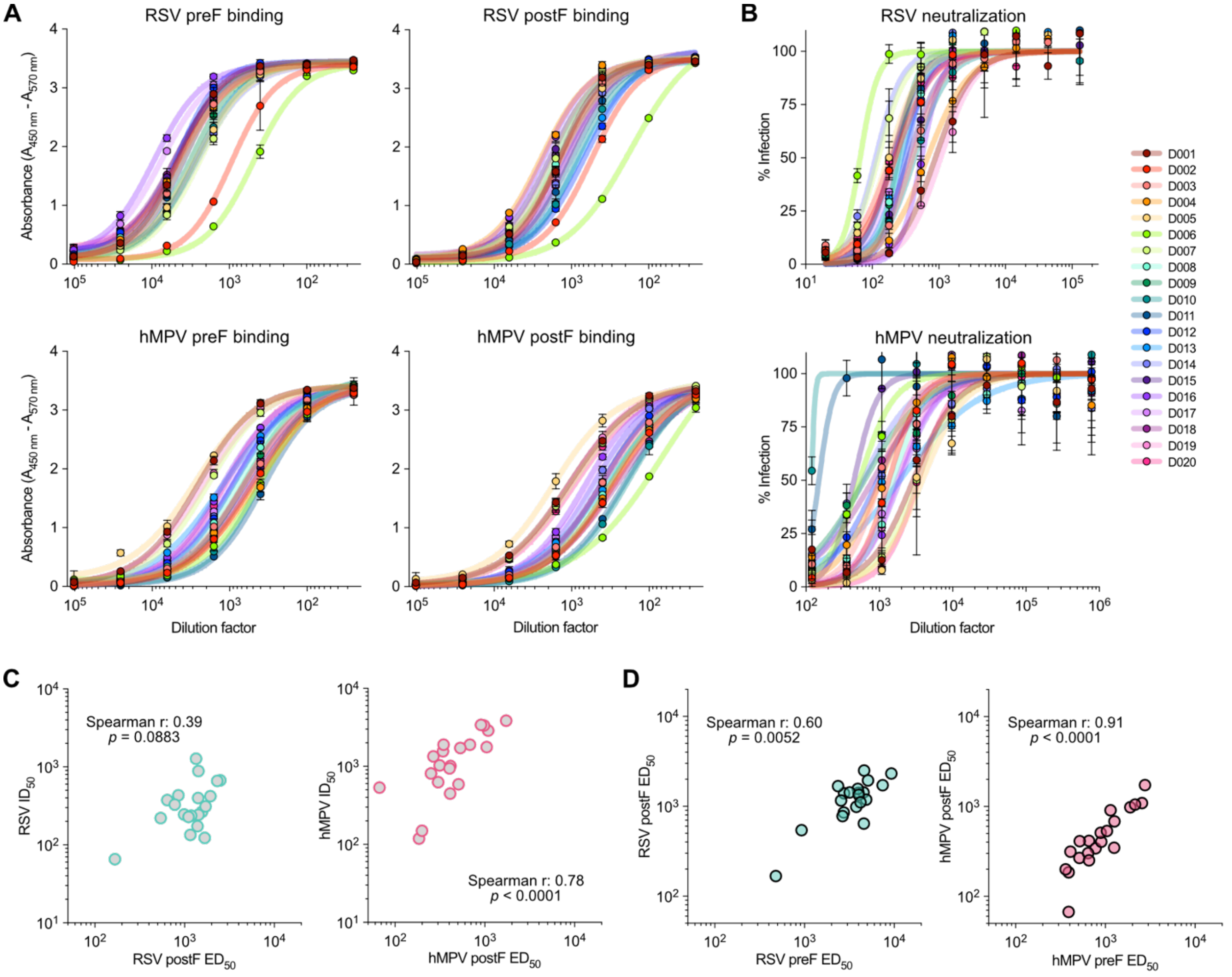


**Figure S2. Circulating antibody responses to RSV and hMPV in healthy adults.**

**A.** ELISA curves for measuring RSV and hMPV preF- and postF-specific IgG binding titers. For each dilution, dots represent the mean absorbance of two replicates and error bars indicate the standard deviation (SD). Curves were fitted using a four-parameter non-linear regression.

**B.** RSV and hMPV neutralization curves. For each dilution, dots represent the mean percentage of infection and error bars indicate the SD. Curves were fitted using a four-parameter non-linear regression.

**C.** Relationship between postF-binding antibody levels and neutralizing titers for RSV and hMPV, analyzed using a non-parametric Spearman correlation.

**D.** Relationship between preF- and postF-binding antibody levels for RSV and hMPV, analyzed using a non-parametric Spearman correlation.

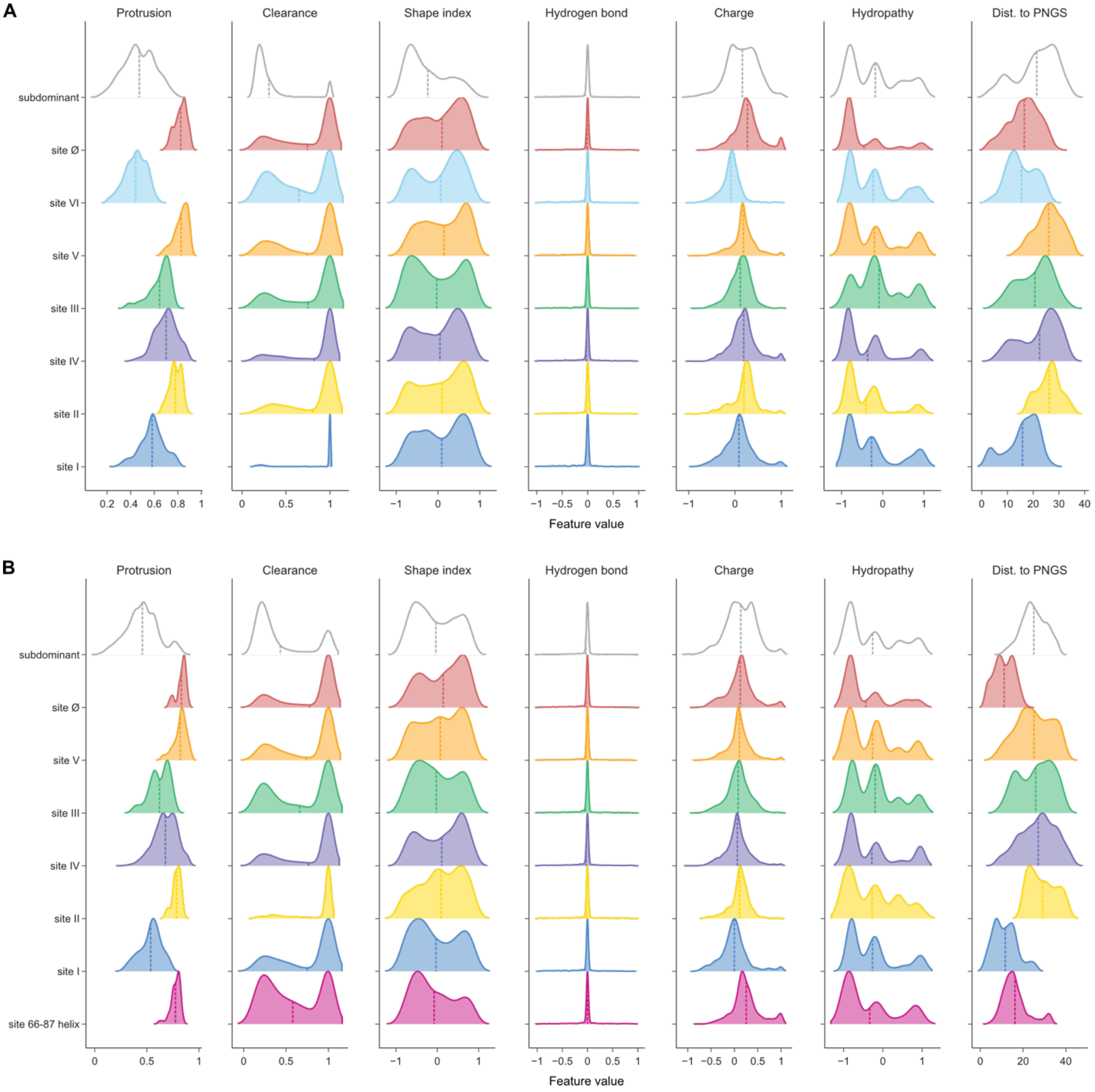


**Figure S3. Antigen surface-centric characterization of pneumovirus preF proteins.**

**A**, **B**. Distribution of geometric, chemical and steric features values among individual surface vertices reported per antigenic site for RSV preF (**A**) and hMPV preF (**B**). Dist., distance; PNGS, potential N-linked glycosylation site.

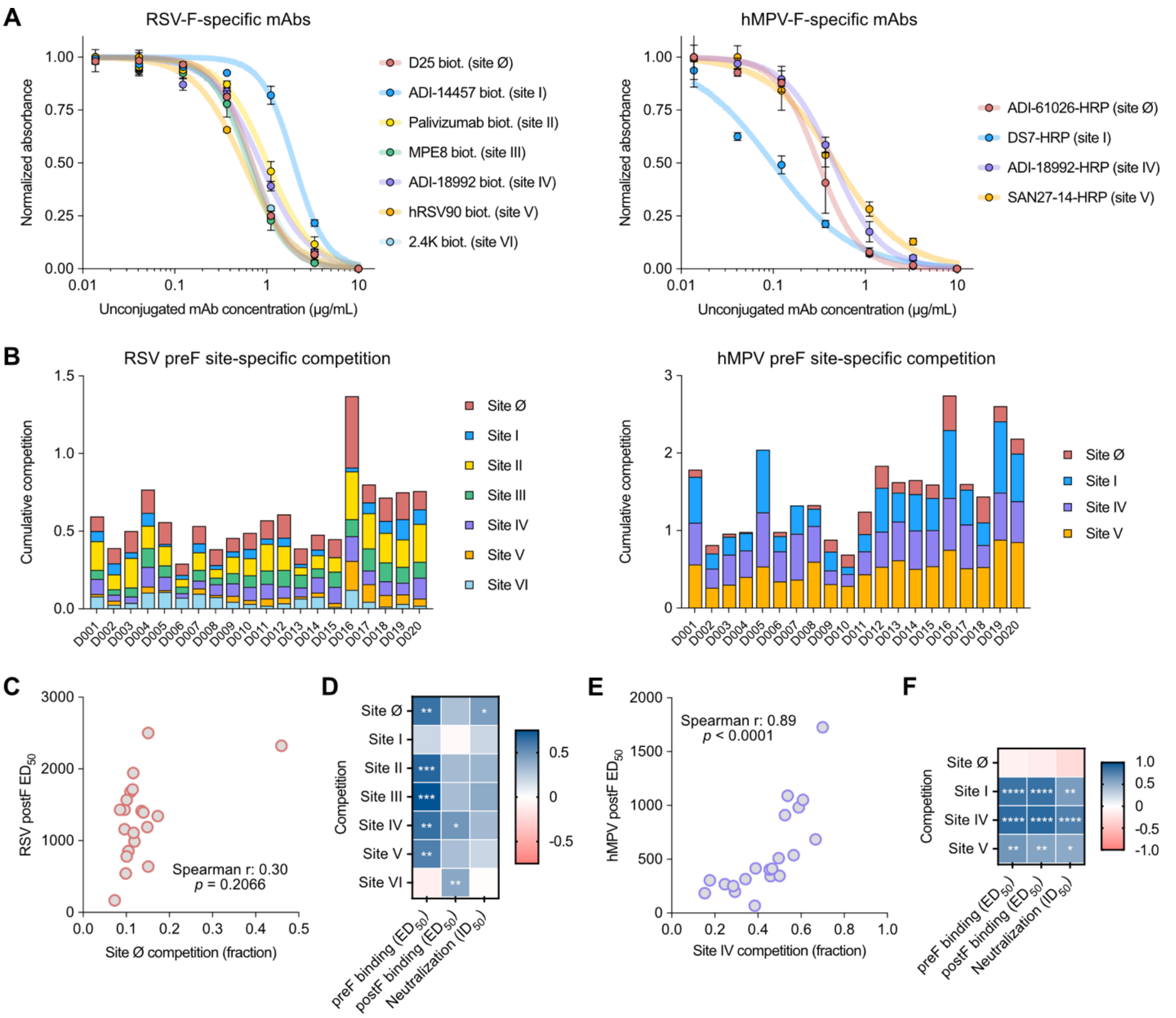


**Figure S4. Pneumovirus-specific memory B cells and circulating antibodies exhibit broad preF specificities.**

**A.** Normalized self-competition ELISA curves (unconjugated vs. biotinylated or HRP-conjugated mAbs) used to determine the site-specific preF binding of circulating antibodies. The level of competition (reported as a fraction) was normalized by defining the signal obtained in the presence of 10 μg/mL of unconjugated mAb as 100% competition (fraction of 1) and the signal without unconjugated mAb as 0% competition (fraction of 0). For each concentration of unconjugated mAb, dots represent the mean normalized absorbance of two replicates and error bars indicate the SD. Curves were fitted using a four-parameter non-linear regression. The assay covered antigenic sites Ø, I, II, III, IV, V and VI for RSV and antigenic sites Ø, I, IV and V for hMPV.

**B.** Cumulative RSV and hMPV preF site-specific competition by circulating antibodies shown per donor.

**C.** Relationship between RSV preF site Ø competition and RSV postF-binding antibody levels, analyzed using a non-parametric Spearman correlation.

**D.** Correlation matrix summarizing the relationships between binding and neutralizing antibody titers with preF site-specific competition for RSV. *$p < 0.05$; ** $p < 0.01$; *** $p < 0.001$; **** $p < 0.0001$.

**E.** Relationship between hMPV preF site IV competition and hMPV postF-binding antibody levels, analyzed using a non-parametric Spearman correlation.

**F.** Correlation matrix summarizing the relationships between binding and neutralizing antibody titers with preF site-specific competition for hMPV. *$p < 0.05$; ** $p < 0.01$; *** $p < 0.001$; **** $p < 0.0001$.

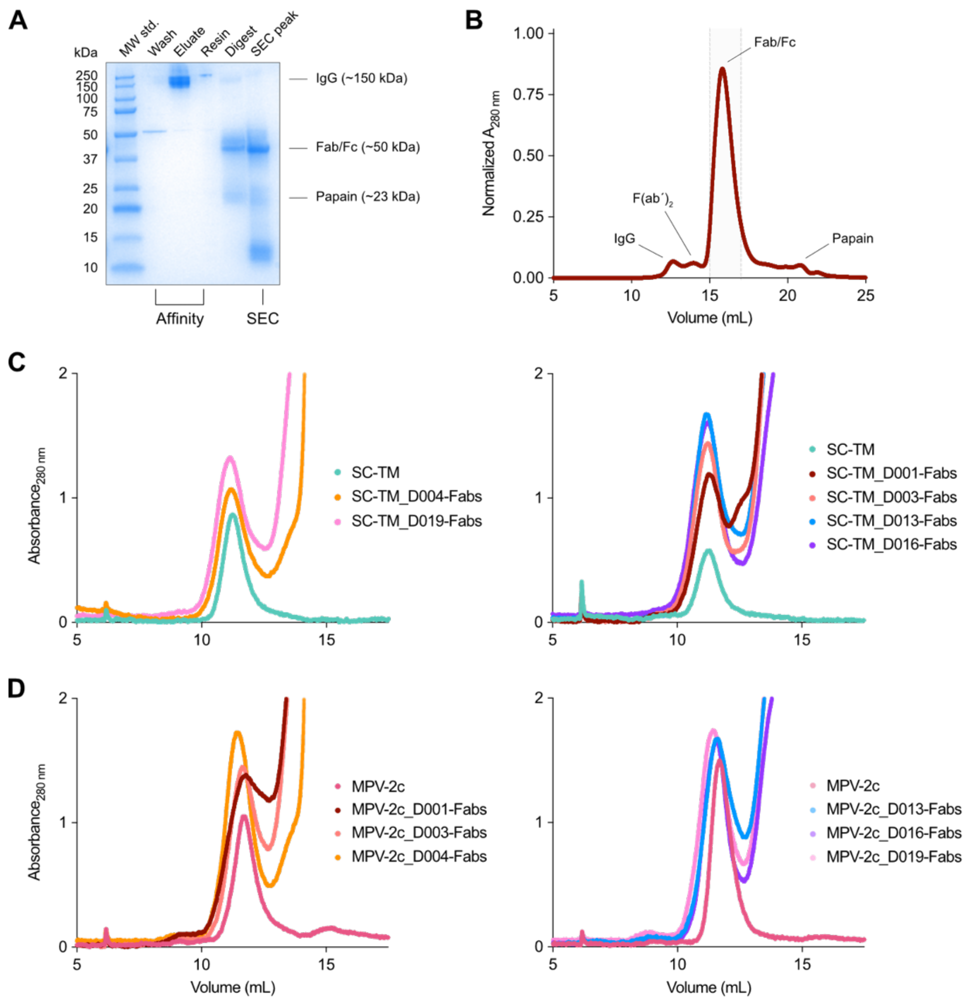


**Figure S5. Isolation of human polyclonal Fabs from plasma and purification of Fab-RSV preF or Fab-hMPV preF immune complexes.**

**A.** Representative SDS-PAGE with Coomassie staining analysis of fractions collected during the isolation of human polyclonal Fabs by CaptureSelect IgG-Fc based affinity chromatography (left) and SEC (right). Gel corresponds to the sample of donor D003.

**B.** Representative SEC trace of CaptureSelect IgG-Fc purified human polyclonal Fabs on a Superdex S200 increase 10/300 GL column. Peak of interest is highlighted in light gray. Trace corresponds to the sample of donor D001.

**C.** SEC traces of SC-TM alone and in complex with RSV preF-targeting polyclonal Fabs.

**D.** SEC traces of MPV-2c alone and in complex with hMPV preF-targeting polyclonal Fabs.

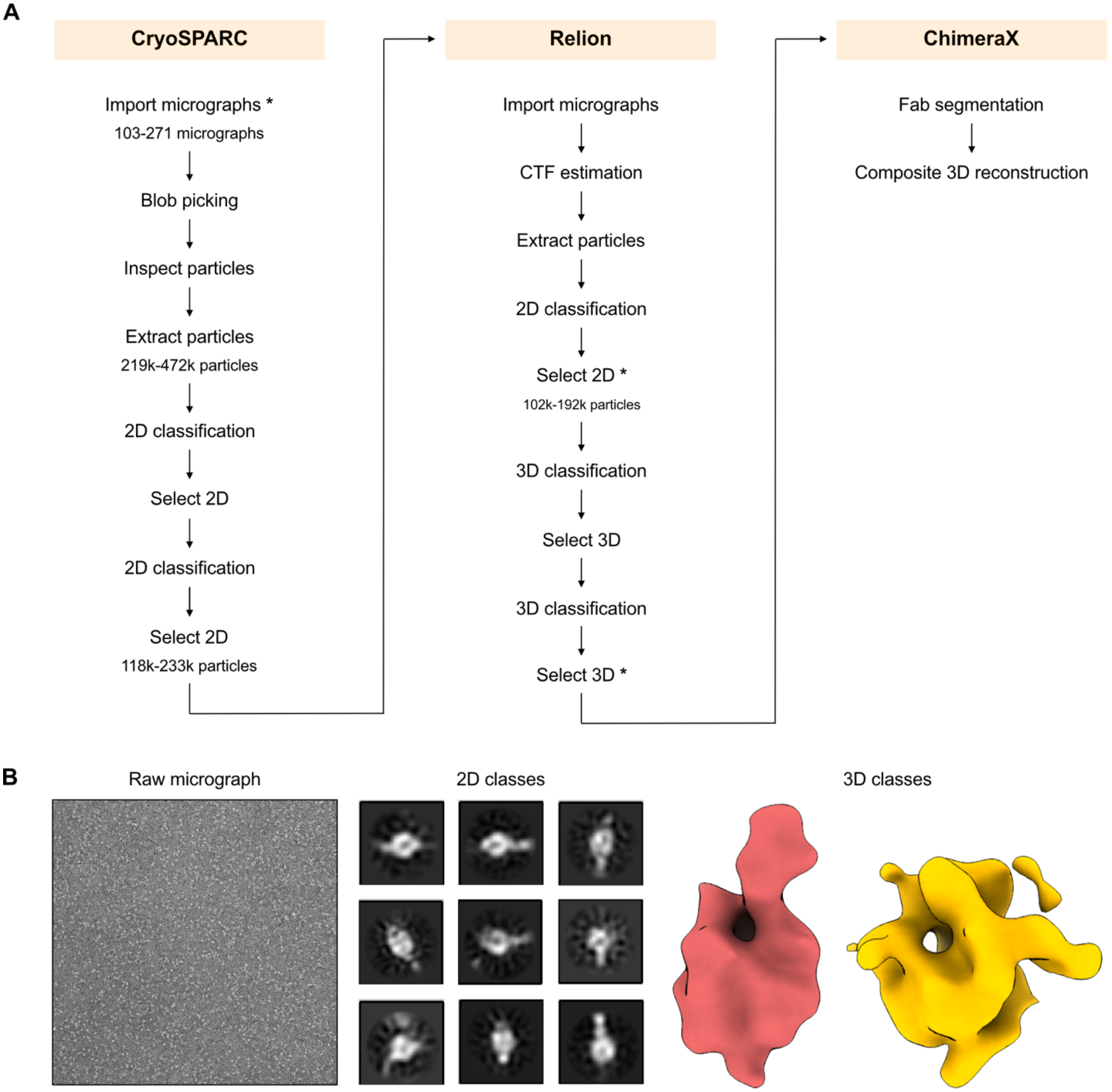


**Figure S6. Negative-stain EMPEM data processing workflow.**

**A.** Representative negative-stain EM data processing workflow using CryoSPARC[4], Relion[5] and ChimeraX[6,7]. Reported numbers indicate the minimum and maximum values observed across all datasets. Steps with an asterisk are visually exemplified in **B**.

**B.** Representative raw micrograph, 2D class averages and 3D reconstructions corresponding to the donor D004 polyclonal Fabs-SC-TM complexes dataset.

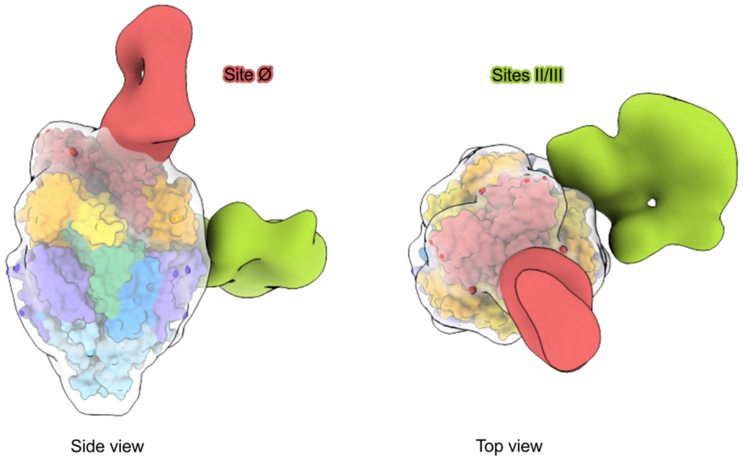


**Figure S7. Epitope mapping strategy.**
Antigenic sites targeted by polyclonal Fabs were identified by visual inspection of the RSV or hMPV preF 3D models (SC-TM[1] or MPV-2c[2], respectively), colored by antigenic site and fitted into preF-Fab 3D reconstructions generated using the negative-stain EMPEM analysis workflow. Side and top views shown correspond to the donor D001 polyclonal Fabs-SC-TM complexes data set.

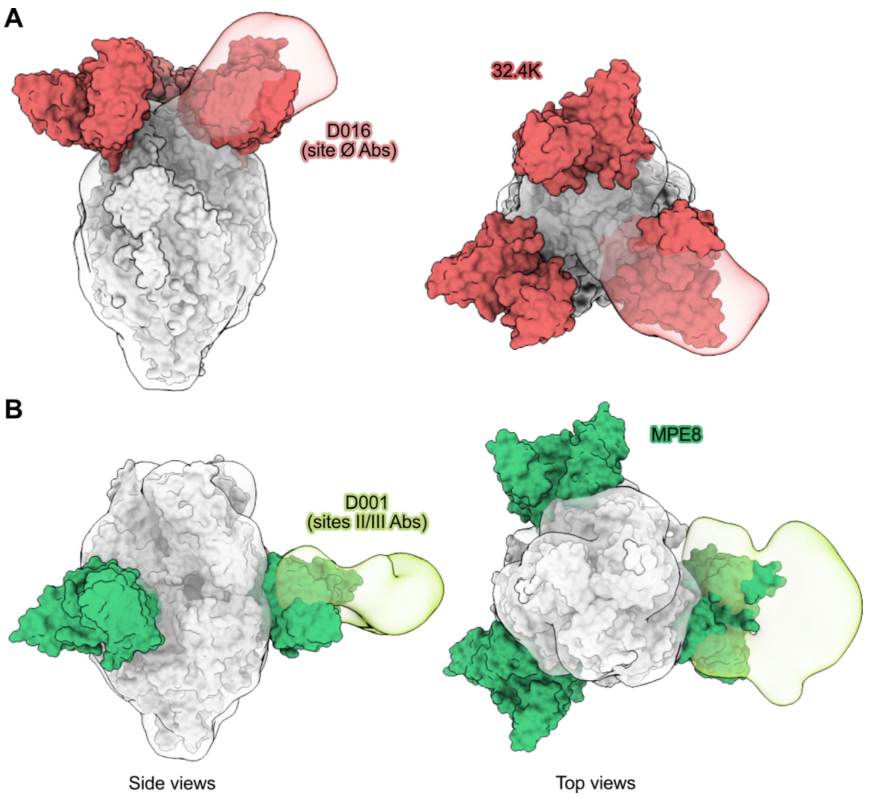


**Figure S8. Resemblance of polyclonal Fab binding modes to public antibody classes.**

**A.** Site Ø-directed polyclonal antibodies from donor D016 targeted the apex epitope similarly to the RSV public antibody class represented by mAb 32.4K[8]. Fitting of the high-resolution structure of the 32.4K Fab-RSV preF complex (PDB: 7LUC) into the polyclonal Fab-preF reconstruction revealed a highly similar binding mode.

**B.** Sites II/III-directed polyclonal antibodies from donor D001 targeted an equatorial epitope similarly to the RSV public antibody class represented by mAb MPE8[9]. Fitting of the high-resolution structure of the MPE8 Fab-RSV preF complex (PDB: 5U68)[8] into the polyclonal Fab-preF reconstruction revealed a highly similar binding mode.

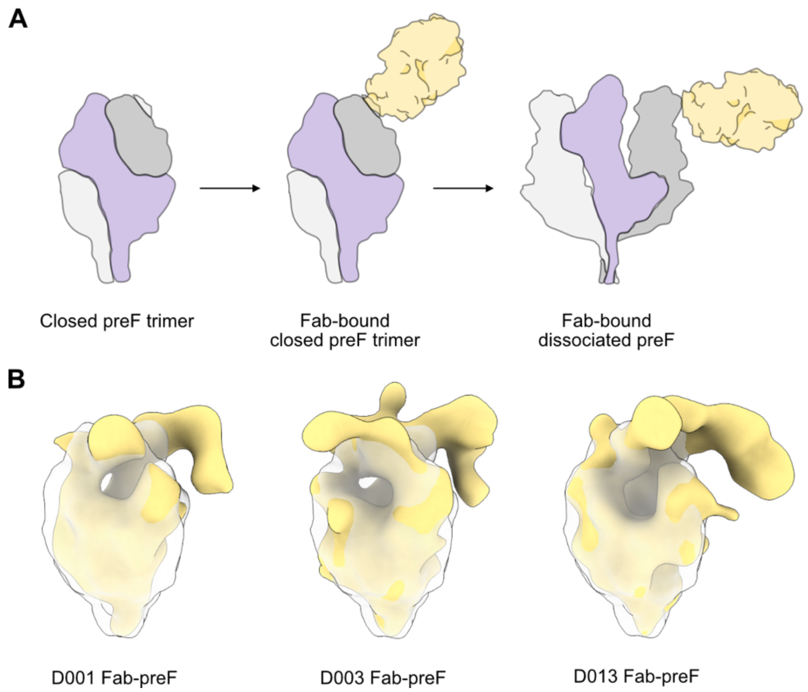


**Figure S9. Antibody-mediated pneumovirus preF trimer destabilization.**

**A.** Schematic illustration of preF trimer destabilization upon antibody binding.

**B.** Representative 3D reconstructions of polyclonal Fabs bound to a partially destabilized pneumovirus preF. 3D reconstructions were fitted into a low-resolution map of the RSV preF trimer.

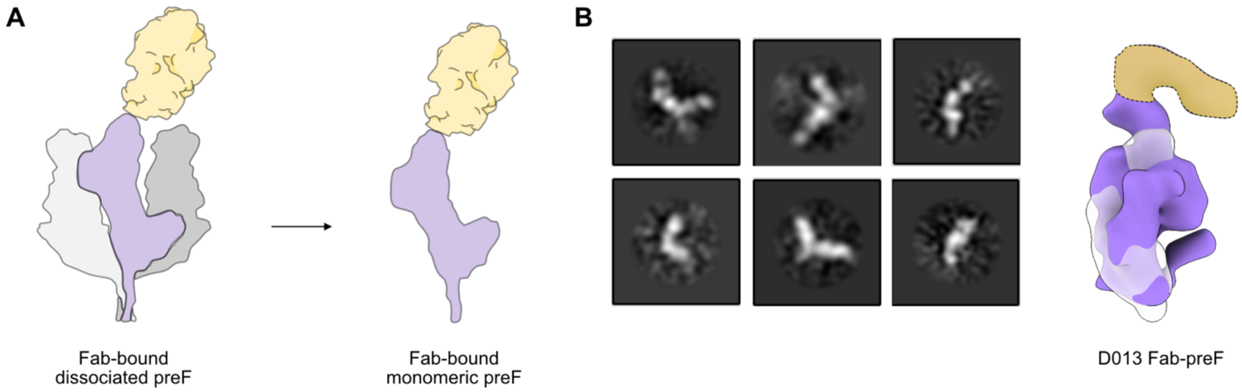


**Figure S10. Monomeric preF-Fab complexes.**

**A.** Schematic illustration of pneumovirus monomeric preF in complex with polyclonal Fabs.

**B.** Representative 2D class averages and 3D reconstruction resembling a pneumovirus preF in its monomeric form in complex with polyclonal Fabs. 3D reconstruction was fitted into a low-resolution map of a pneumovirus preF monomer.

**Table S1. EMDB accession codes for refined 3D reconstructions of polyclonal Fabs from healthy adult donors in complex with RSV or hMPV preF.**

| Donor ID | Antigen | Targeted epitope | EMDB Accession Code |
|---|---|---|---|
| D001 | RSV preF | Site Ø | EMD-100673 |
| D001 | RSV preF | Sites II/III | EMD-100674 |
| D003 | RSV preF | Site Ø | EMD-100675 |
| D003 | RSV preF | Sites IV/V | EMD-100676 |
| D004 | RSV preF | Site Ø | EMD-100679 |
| D004 | RSV preF | Sites II/V | EMD-100681 |
| D013 | RSV preF | Site Ø | EMD-100682 |
| D016 | RSV preF | Site Ø | EMD-100683 |
| D016 | RSV preF | Sites IV/V | EMD-100684 |
| D019 | RSV preF | Site Ø | EMD-100686 |
| D001 | hMPV preF | Site Ø | EMD-100689 |
| D001 | hMPV preF | Site V | EMD-100691 |
| D001 | hMPV preF | Sites II/V | EMD-100692 |
| D003 | hMPV preF | Site Ø | EMD-100693 |
| D004 | hMPV preF | Site Ø | EMD-100694 |
| D019 | hMPV preF | Site Ø | EMD-100695 |
| D019 | hMPV preF | Sites II/V | EMD-100696 |
